# Key features of inhibitor binding to the human mitochondrial pyruvate carrier hetero-dimer

**DOI:** 10.1101/2021.12.14.472537

**Authors:** Sotiria Tavoulari, Tom J.J. Schirris, Vasiliki Mavridou, Chancievan Thangaratnarajah, Martin S. King, Daniel T.D. Jones, Shujing Ding, Ian M. Fearnley, Edmund R.S. Kunji

**Author notes:** **Corresponding authors:** Sotiria Tavoulari, Medical Research Council Mitochondrial Biology Unit, University of Cambridge, Keith Peters Building, Cambridge Biomedical Campus, Hills Road, Cambridge, CB2 0XY, United Kingdom, Edmund R.S. Kunji, Medical Research Council Mitochondrial Biology Unit, University of Cambridge, Keith Peters Building, Cambridge Biomedical Campus, Hills Road, Cambridge, CB2 0XY, United Kingdom. Department of Pharmacology and Toxicology, Radboud Institute for Molecular Life Sciences, Radboud Center for Mitochondrial Medicine, Radboud University Medical Center, Nijmegen, 6500 HB, The Netherlands. Groningen Biomolecular Sciences and Biotechnology Institute, Membrane Enzymology, University of Groningen, Nijenborgh 4, 9747 AG, Groningen, The Netherlands. Department of Biological Chemistry and Molecular Pharmacology, Harvard Medical School, LHRRB 201, 45 Shattuck St., 02115, Boston, MA, USA.

## Abstract

The mitochondrial pyruvate carrier (MPC) has emerged as a promising drug target for metabolic disorders, including non-alcoholic steatohepatitis and diabetes, metabolically dependent cancers and neurodegenerative diseases. Human MPC is a protein complex, but the composition of its active form is debated and the mechanisms of transport and inhibition are not resolved. We have recombinantly expressed and purified the human hetero-complex MPC1L/MPC2 and demonstrate that it is a functional hetero-dimer, like the yeast MPC hetero-dimers. Unlike the latter, human MPC1L/MPC2 binds the known inhibitors with high potencies. We identify the essential chemical features shared between these structurally diverse inhibitors and demonstrate that high affinity binding is not attributed to covalent bond formation with MPC cysteines, as previously thought. We also identify 14 new inhibitors of MPC, one outperforming the most potent compound UK5099 by tenfold. Two of them are the commonly prescribed drugs entacapone and nitrofurantoin, suggesting possible off-target mechanisms associated with their adverse effects. This work advances our understanding of MPC inhibition and will accelerate the development of clinically relevant MPC modulators.

## INTRODUCTION

The mitochondrial pyruvate carrier (MPC) is responsible for the import of pyruvate, produced during glycolysis, into the mitochondrial matrix where it enters the TCA cycle after its conversion to acetyl-CoA. This transport step links the cytosolic and mitochondrial energy metabolism, providing MPC with a key role in the metabolic fate of the cell. In addition, MPC has a pivotal role in several central metabolic pathways, such as gluconeogenesis, fatty acid and amino acid metabolism. Consequently, MPC is being investigated as a promising modulator of metabolic disorders, such as type-2 diabetes, non-alcoholic fatty liver disease (NAFLD) and non-alcoholic steatohepatitis (NASH) (Colca *et al*., 2018, Gray *et al*., 2015, Harrison *et al*., 2020, McCommis *et al*., 2017). It has also been proposed as a target for the treatment of neurodegenerative diseases (Divakaruni *et al*., 2017, Ghosh *et al*., 2016, Quansah *et al*., 2018) and metabolically dependent cancers (Bader *et al*., 2019, Corbet *et al*., 2018, Elia *et al*., 2019, Tompkins *et al*., 2019).

MPC is a protein complex formed by two small homologous membrane proteins (Bricker *et al*., 2012, Herzig *et al*., 2012). In yeast, two alternative hetero-complexes form depending on the carbon source availability; Mpc1/Mpc2 in fermentative conditions and Mpc1/Mpc3 in respiratory conditions (Bender *et al*., 2015). We have recently reported the purification and functional reconstitution of the yeast MPCs and have demonstrated that the functional unit is a hetero-dimer (Tavoulari *et al*., 2019). We have also shown that the individual yeast proteins are able to form homo-dimers in the absence of the other protomer, but they are inactive (Tavoulari *et al*., 2019). In humans, two proteins, MPC1 and MPC2, are ubiquitously expressed to form the MPC1/MPC2 complex, present in all tissues (Vanderperre *et al*., 2016). An additional complex can be formed in the testis, specifically in post meiotic germ cells (Vanderperre *et al*., 2016). This complex is formed by the same MPC2 protomer and MPC1L (Vanderperre *et al*., 2016), which is homologous to MPC1 (**Figure EV1**). In separate studies, the individual human MPC2 protein has been proposed to be active for pyruvate transport as an autonomous entity (Nagampalli *et al*., 2018) and the individual MPC1 and MPC2 protomers have been proposed to bind small molecule inhibitors in the absence of their partner protein (Lee *et al*., 2020), suggesting fundamental differences in MPC functional complexes between yeast and humans. However, human MPC hetero-complexes have not yet been reconstituted into liposomes in a functional state, and their properties have not been determined or compared with homo-complexes.

A few different, often clinically relevant compounds, have been proposed to inhibit pyruvate-driven respiration or pyruvate transport in isolated mitochondria, functions attributed to MPC. The best characterized compounds for monitoring MPC inhibition are α-cyano-cinnamates and derivatives, such as the α-cyano-4-hydroxy-cinnamate (CHC) (Halestrap & Denton, 1974, Halestrap & Denton, 1975), or more potent analogues, such as the α-cyano-β-(1-phenylindol-3-yl)-acrylate, also known as UK5099 (Halestrap, 1975). Other structurally diverse compounds have been proposed more recently. Zaprinast, an inhibitor of cGMP-specific phosphodiesterase (PDE) and lead compound used for the development of sildenafil (Viagra) inhibits pyruvate-driven oxygen consumption in brain mitochondria and pyruvate uptake in liver mitochondria (Du *et al*., 2013). Silibinin, an adjunctive therapy in chronic hepatitis and cirrhosis and valproic acid, a drug used to treat epilepsy, bipolar disorder and migraine headaches have also been implicated in pyruvate uptake inhibition (Benavides *et al*., 1982, Colturato *et al*., 2012). The anti-cancer agent lonidamine has been shown to inhibit pyruvate uptake in isolated mitochondria and inhibits the monocarboxylate transporters (MCTs) with lower affinities (Nancolas *et al*., 2016, Nath *et al*., 2016). Importantly, MPC is a newly identified target for known insulin sensitizers, called thiazolidinediones (TZDs) (Colca *et al*., 2013, Divakaruni *et al*., 2013), also known to exert their action via the peroxisome proliferator-activated receptor gamma (PPARγ) (Nanjan *et al*., 2018, Soccio *et al*., 2014). New generation TZDs, including MSDC-0160 (mitoglitazone) and MSDC-0602K, with decreased affinity for PPARγ, have been developed (Colca *et al*., 2014) and are being considered in clinical trials for the treatment of non-alcoholic steatohepatitis (Harrison *et al*., 2020) and Alzheimer’s disease (Shah *et al*., 2014), emphasizing the importance of understanding the molecular nature of MPC inhibition.

In this study, we have established *in vitro* systems relying on recombinant, heterologously expressed human MPC proteins to investigate the key features of inhibitor binding. First, we have shown that the human MPC is a hetero-dimeric complex mediating pyruvate transport in proteoliposomes. Next, we have compared a range of structurally diverse compounds for binding and inhibitory potency and have identified the critical and shared chemical properties important for MPC inhibition, whilst new inhibitors with higher potency have been identified.

## RESULTS

### Pyruvate transport and ligand binding by human MPC hetero-dimers

Human MPC hetero-complexes MPC1/MPC2 and MPC1L/MPC2 were expressed in the *S. cerevisiae mpc* triple deletion strain SHY15 (Herzig *et al*., 2012) and purified by nickel-affinity chromatography using a cleavable eight-histidine tag at the C-terminus of MPC2, while MPC1 or MPC1L remained untagged. From the two complexes tested, only MPC1L/MPC2 was purified with good yields (>0.7 mg of protein per gram of isolated mitochondria) and was characterized by an equimolar composition of the two protomers (**Figure 1a, left**), similar to the yeast Mpc1/Mpc3 hetero-dimer (Tavoulari *et al*., 2019). The size of the human MPC1L/MPC2 was also similar to the yeast Mpc1/Mpc3 (Tavoulari *et al*., 2019) by size-exclusion chromatography in the same detergent/lipid mix (**Figure EV2a**), suggesting that it also forms a hetero-dimer. The calculated mass by size-exclusion chromatography linked to multi-angle laser light scattering (SEC-MALLS) (43 ± 6 kDa) along with the equimolar presence of the two protomers were also consistent with the formation of a hetero-dimer (**Figure EV2b**).

**Figure 1.**
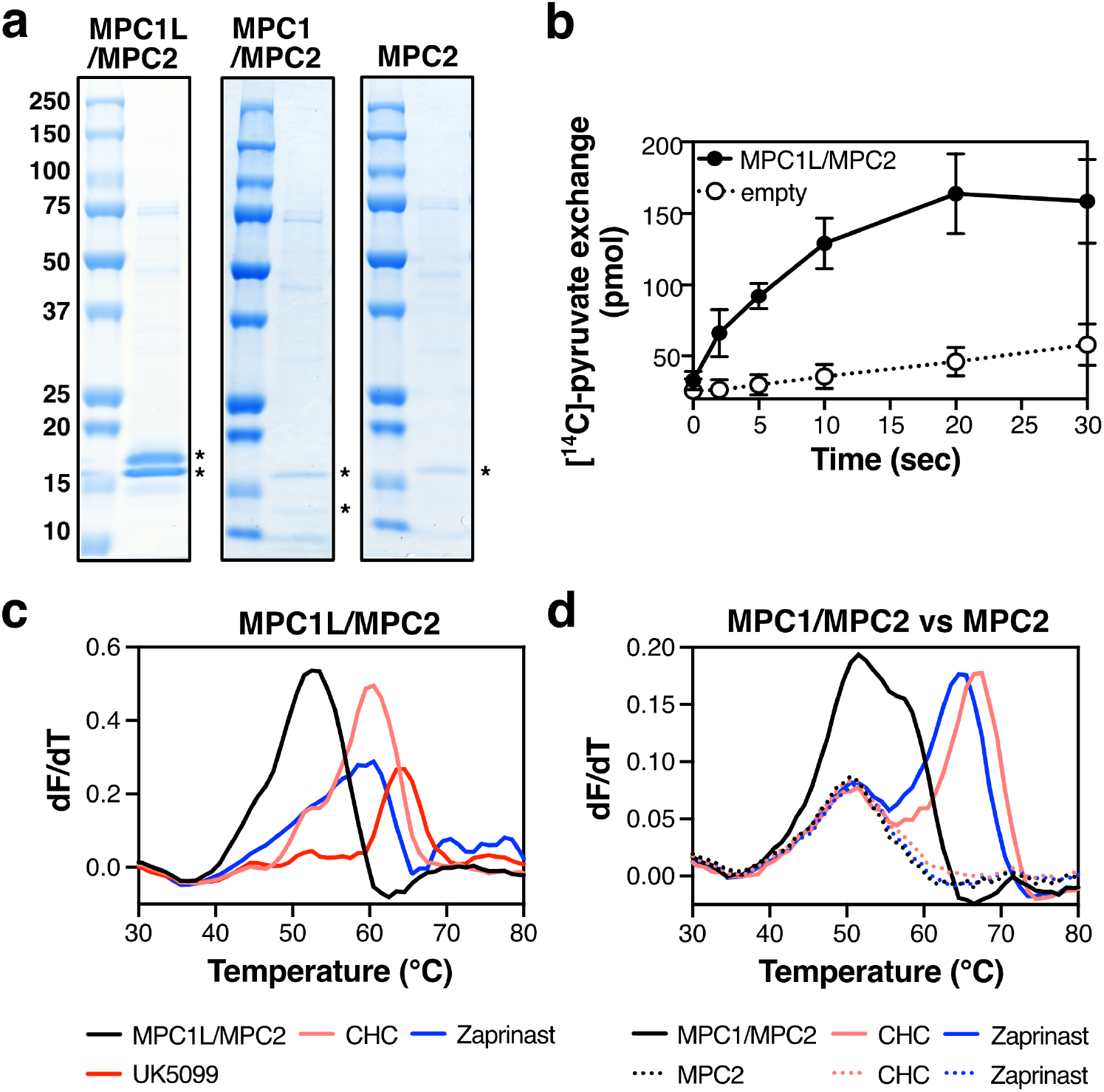
Pyruvate transport and ligand binding by human MPC. (**a**) Νickel affinity-purified MPC1L/MPC2 hetero-dimer (left), MPC1/MPC2 hetero-dimer (middle) and MPC2 protomer (right) analyzed by SDS-PAGE and visualized by Coomassie Blue stain. MPC proteins are indicated by asterisks. (**b**) Time course of pyruvate homo-exchange at a ΔpH of 1.6 for the MPC1L/MPC2 hetero-dimer reconstituted into liposomes. (**c-d**) CPM thermostability shift assays for MPC hetero-dimers and individual MPC2 in the presence of absence of 100 μM MPC inhibitors. Data information: In (**b**), data represent the mean ± s.d. of six biological repeats, each performed with two technical replicates. In (**c**-**d)**, data are representative of two biological repeats, each performed with three or two technical replicates, respectively.

We then assessed the ability of the MPC1L/MPC2 dimer to transport pyruvate when reconstituted into liposomes (**Figure 1b**), as used for the yeast MPC (Tavoulari *et al*., 2019). The initial uptake rate of pyruvate transport was 0.87 ± 0.17 µmol/min/mg protein, similar to the rate previously measured for the yeast Mpc1/Mpc3 (Tavoulari *et al*., 2019). To evaluate ligand binding we used thermostability shift assays. We monitored the unfolding of the protein population in a temperature ramp in the presence of 7-diethylamino-3-(4-maleimidophenyl)-4-methylcoumarin (CPM), which reacts with buried cysteine residues that become exposed due to denaturation (**Figure 1c and d**). In the presence of ligand, the protein population shifts to higher melting temperatures, due to the increased number of interactions forming (Crichton *et al*., 2015). A decrease in thermostability is also possible, if a compound destabilizes the complex. Unlike what we had observed previously for the yeast protein (Tavoulari *et al*., 2019), the thermostability results for the MPC1L/MPC2 hetero-dimer (**Figure 1c**) showed strong shifts in the presence of different MPC inhibitors. Specifically, the melting temperature of the unliganded MPC1L/MPC2 shifted from 51.3 ± 0.2 °C to 59.6 ± 0.4 °C, 58.8 ± 0.4 °C and 62.5 ± 0.9 °C in the presence for the previously known inhibitors CHC, zaprinast, and UK5099, respectively (**Figure 1c** and **Supplementary Table EV1**). Overall, these results support a hetero-dimeric functional unit for the human complex, with high affinity for inhibitors. We also purified the MPC1/MPC2 complex, albeit with very low yields (<0.1 mg of protein per gram of isolated crude mitochondria) and a ratio of MPC1 to MPC2 of approximately 1:2, possibly suggesting excess of MPC2 during co-expression and/or co-purification (**Figure 1a, middle**). In thermostability shift assays for MPC1/MPC2, we observed strong shifts in the presence of CHC and Zaprinast, consistent with high affinity inhibitor binding also in this complex. However, both in the unliganded and liganded forms the protein presented two main peaks, one of which did not shift in the temperature ramp. We then attempted to express and purify the protomers of this complex individually to dissect the unfolding profile of MPC1/MPC2. Although no MPC1 protein could be purified alone, we were able to purify MPC2 alone, a protein previously proposed to form functional homo-complexes (Nagampalli *et al*., 2018). The individually purified MPC2 was only produced in very low yields (**Figure 1a, right**) and could not be tested for transport activity. We were, however, able to test it for inhibitor binding in thermostability shift assays, where neither CHC or zaprinast, showed any shift (**Figure 1d**). Additionally, the MPC2 unfolding curve aligned with the first peak of MPC1/MPC2, which does not shift in the presence of inhibitors. From these results, we conclude that binding of inhibitors to MPC2 homomers, also included in the MPC1/MPC2 sample, is absent or very weak.

### Shared chemical features enabling MPC inhibition by structurally distinct molecules

A variety of small molecules (**Figure 2a**) have been proposed to alter pyruvate-driven mitochondrial metabolism or inhibit pyruvate transport. Some of the compounds belong to the same class, such as the cyano-cinnamates (*e.g*., UK5099 and CHC) and the thiazolidinediones (*e.g*., rosiglitazone, pioglitazone and mitoglitazone), but overall, the MPC inhibitors are structurally diverse. To understand inhibitor binding, we performed the first comparative analysis of inhibitory potency using functionally reconstituted purified MPC.

**Figure 2.**
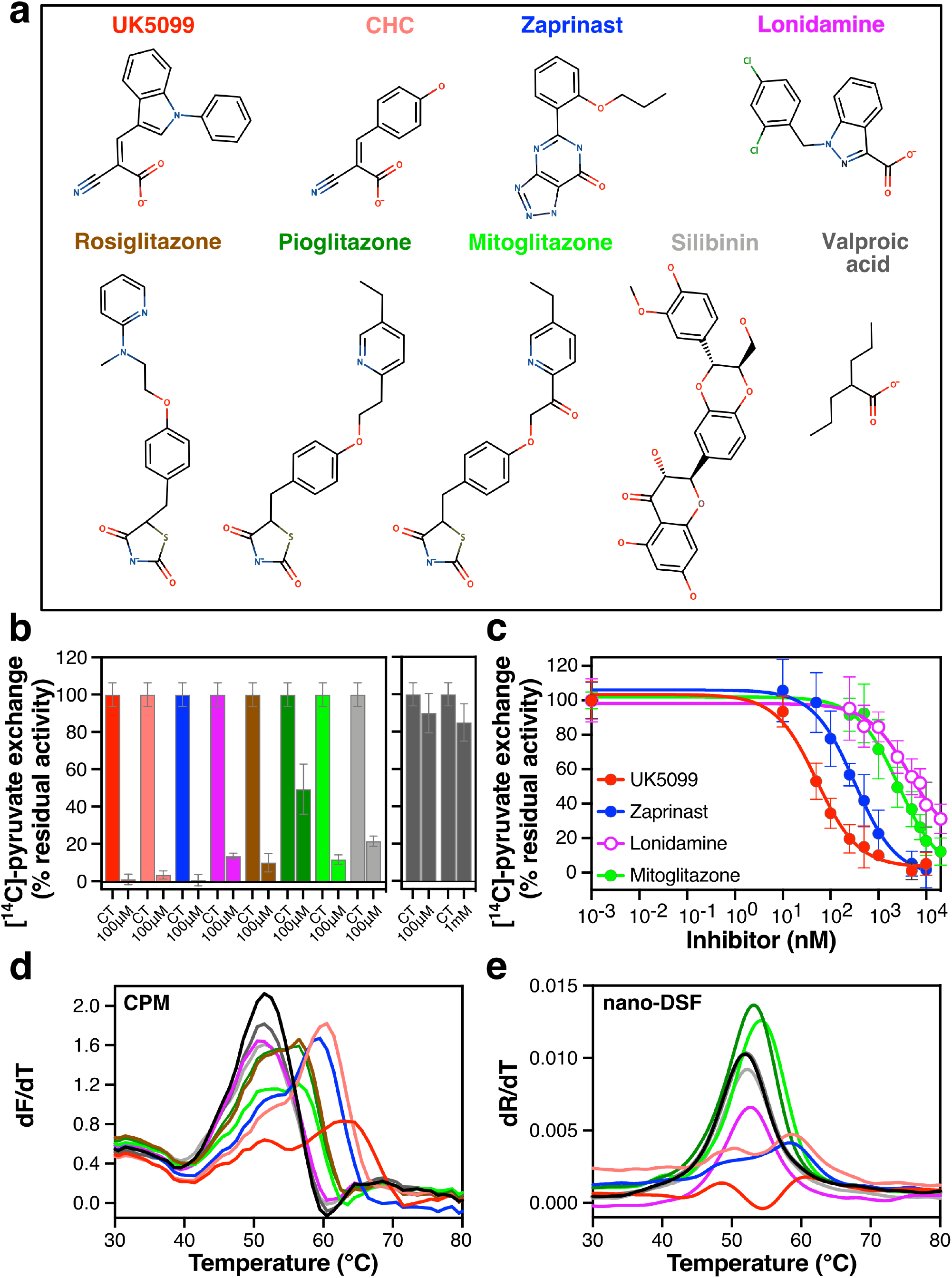
Comparative analyses of ligand binding and pyruvate transport inhibition by structurally distinct small molecules. (**a**) Chemical structures of previously claimed MPC inhibitors. The color coding is consistently used in all panels. (**b**) The compounds listed in (**a**) were tested for inhibition of pyruvate transport on MPC1L/MPC2 proteoliposomes at 100 μM and the remaining transport activity was compared to that in their absence (CT). (**c**) Inhibition of [^14^C]-pyruvate homo-exchange by UK5099 (10–10,000 nM), zaprinast (10– 10,000 nM), lonidamine (250–20,000 nM) and mitoglitazone (250–20,000 nM). (**d**) First derivative of MPC1L/MPC2 unfolding profiles with or without compounds (100 μM) via the CPM thermostability assay. (**e**) First derivative of MPC1L/MPC2 unfolding profiles (330 nm/350 nm ratio) with or without compounds (100 μM) via nano-DSF. Data information: In (**b**), data points represent the mean ± s.d. of two biological repeats performed in triplicate. In (**c**), data points represent the mean ± s.d of four (UK5099) or three (zaprinast, lonidamine, mitoglitazone) biological repeats, each performed in triplicate. Results in (**d**-**e**) represent characteristic experiments repeated independently. Averaged data from different biological repeats are summarized in **Table EV1**.

First, we tested all of the listed molecules for their ability to inhibit pyruvate transport by MPC1L/MPC2 at a high concentration of 100 μM (**Figure 2b**). UK5099, CHC and zaprinast, completely inhibited pyruvate transport at this concentration, whereas lonidamine, the TZDs and silibinin inhibited to a lower extent, ranging from 10 to 50 % of residual activity compared to control conditions. In contrast, the effect of valproic acid on pyruvate transport was minimal even at 1 mM final concentration (**Figure 2b**, right panel). To confirm that UK5099, CHC and zaprinast have a higher inhibitory potency than the TZDs, lonidamine or silibinin, we determined IC_50_-values of pyruvate transport inhibition for prototypic MPC inhibitors, as shown in **Figure 2c**. Indeed, UK5099 showed the highest inhibitory potency followed by zaprinast, with IC_50_-values of 52.6 ± 8.3 nM and 321 ± 42 nM, respectively. In contrast, lonidamine and mitoglitazone had lower potencies, with IC_50_-values in the low micromolar range (4.6 ± 1.3 μM and 2.7 ± 0.8 μM, respectively). Interestingly, UK5099 inhibits human MPC1L/MPC2 with two hundred times higher potency compared to the yeast protein. Zaprinast, lonidamine and the TZDs also have much higher potencies for the human protein (Tavoulari *et al*., 2019), suggesting key differences in the binding site between the two species.

Despite exceptional reliability, the pyruvate transport assay did not constitute a high throughput solution for compound screening. Therefore, we evaluated thermostability shift as a primary screening approach for MPC binders. We tested all compounds listed in **Figure 2a** for their effect on MPC thermostability using two thermostability shift assays; the CPM assay (**Figure 2d**) and nano-differential scanning fluorimetry (nano-DSF), which relies on the intrinsic fluorescence of tryptophan and tyrosine (**Figure 2e**). Both assays yielded similar results for unfolding of the unliganded dimer with apparent melting temperature (Tm) values at 51.3 ± 0.2 °C and 51.9 ± 0.1 °C by the CPM and nano-DSF assay, respectively. Incubation of the dimer with each inhibitor at the same concentration of 100 μM resulted in a range of thermal shifts (**Table EV1**). In the CPM assay, UK5099 yielded the highest thermal shift of all compounds (11.2 ±1.1 °C compared to the no-ligand control), followed by CHC (8.3 ± 0.6 °C) and zaprinast (7.5 ± 0.6 °C), consistent with the transport assay data showing highest inhibitory potencies for these compounds. The TZDs yielded a more moderate shift (up to 5 °C) and valproic acid had no effect at all, also consistent with transport. No thermal shifts were observed for lonidamine and silibinin at a concentration of 100 μM in the CPM assay (**Figure 2d** and **Table EV1**) or by nano-DSF (**Figure 2e** and **Table EV1)**, despite inhibiting pyruvate transport at the same concentration. However, at 1 mM, a positive thermal shift in the CPM assay was observed for silibinin and a negative thermal shift for lonidamine (**Figure EV3**).

Comparisons with the CPM assay showed that a few compounds suffered from fluorescence quenching in nano-DSF. UK5099, CHC and zaprinast shifted the denaturation curve from 51.9 ± 0.1 °C to 61.1 ± 0.6 °C, 58.9 ± 0.4 °C and 58.2 ± 0.3 °C, respectively (**Figure 2e**). Pioglitazone and mitoglitazone resulted in smaller thermal shifts compared to the CPM analysis, reaching Tm values of 53.0 ± 0.2 °C and 53.8 ± 0.4 °C, respectively, and rosiglitazone could not be tested due to autofluorescence. Due to these interferences, the CPM thermostability shift assay was chosen for subsequent investigations of the binding properties. However, data from both CPM and nano-DSF were consistent with UK5099, CHC and zaprinast being the strongest binders.

To understand binding of structurally diverse molecules on MPC, we compared the macromolecular interaction characteristics of the prototypic MPC inhibitors UK5099, zaprinast, lonidamine and mitoglitazone and of the substrate pyruvate. A pharmacophore model was generated by structural alignment, which resulted in a representation of all shared pharmacophore features (**Figure EV4a and g**). Interestingly, all inhibitors share three closely spaced (< 2.95 Å) hydrogen bond acceptor groups (**Figure EV4c-f and g**, groups III, IV, V), also present in the substrate pyruvate (**Figure EV4a, b and g**), suggesting that the inhibitors bind into the MPC substrate-binding pocket in a similar way as pyruvate. All prototypic inhibitors additionally share a central hydrophobic aromatic ring (**Figure EV4c-f and g**, groups I and II), which is not present in pyruvate. Another pharmacophore feature shared by some, but not all inhibitors, consists of a negatively charged ionizable group (*i.e.,* in lonidamine, UK5099, and mitoglitazone) (**Figure EV4b-g**, group VI). However, the presence of this particular feature did not cluster with higher or lower affinity inhibitors.

Overall, all the inhibitors share pyruvate-like hydrogen bond acceptor properties followed by an adjacent central hydrophobic aromatic ring. These combined features are likely to be a minimal requirement for inhibitory potency in the micromolar range and below.

### Compound evolution increases the inhibitory potency and highlights the key chemical features

Next, we aimed to understand and enhance the inhibitory potency using the shared pharmacophore features (**Figure EV4**) of the known MPC inhibitors as a starting point. Because UK5099 showed the highest inhibitory potency (**Figure 2**), while sharing the three-hydrogen bond acceptors (**Figure EV4g**), we chose the carboxyl and cyanide functional groups of UK5099 as a molecular template to search for new chemical entities. We explored over 1,120,000 small molecules (*i.e.,* Chembridge Express and Core libraries) for compounds that contain this molecular template as well as an adjacent central aromatic ring.

Compound 1, (**Figure 3a**), which embodies all the main features of our pharmacophore, was a good MPC binder, as it yielded a thermostability shift to 60.4 ± 1.0 °C from 51.3 ± 0.6 °C in the absence of the ligand (**Figure 3c** and **d** and **Table EV1**). This result demonstrated the validity of the pharmacophore model (**Figure EV4g**) and the importance of these features for MPC binding. An extension in the region following the central aromatic ring moiety (compound 2, **Figure 3a**), with an additional aromatic ring, increased the thermostability to 63.4 ± 0.9 °C, more than compound 1 or UK5099 (**Figure 3c** and **d** and **Table EV1**). Indeed, as demonstrated by pyruvate transport inhibition in proteoliposomes, compound 2 is a more potent inhibitor than UK5099, with an IC_50_-value at 12.4 ± 4.6 nM, compared to 52.6 ± 8.3 nM for UK5099 (**Figure 3e**).

**Figure 3.**
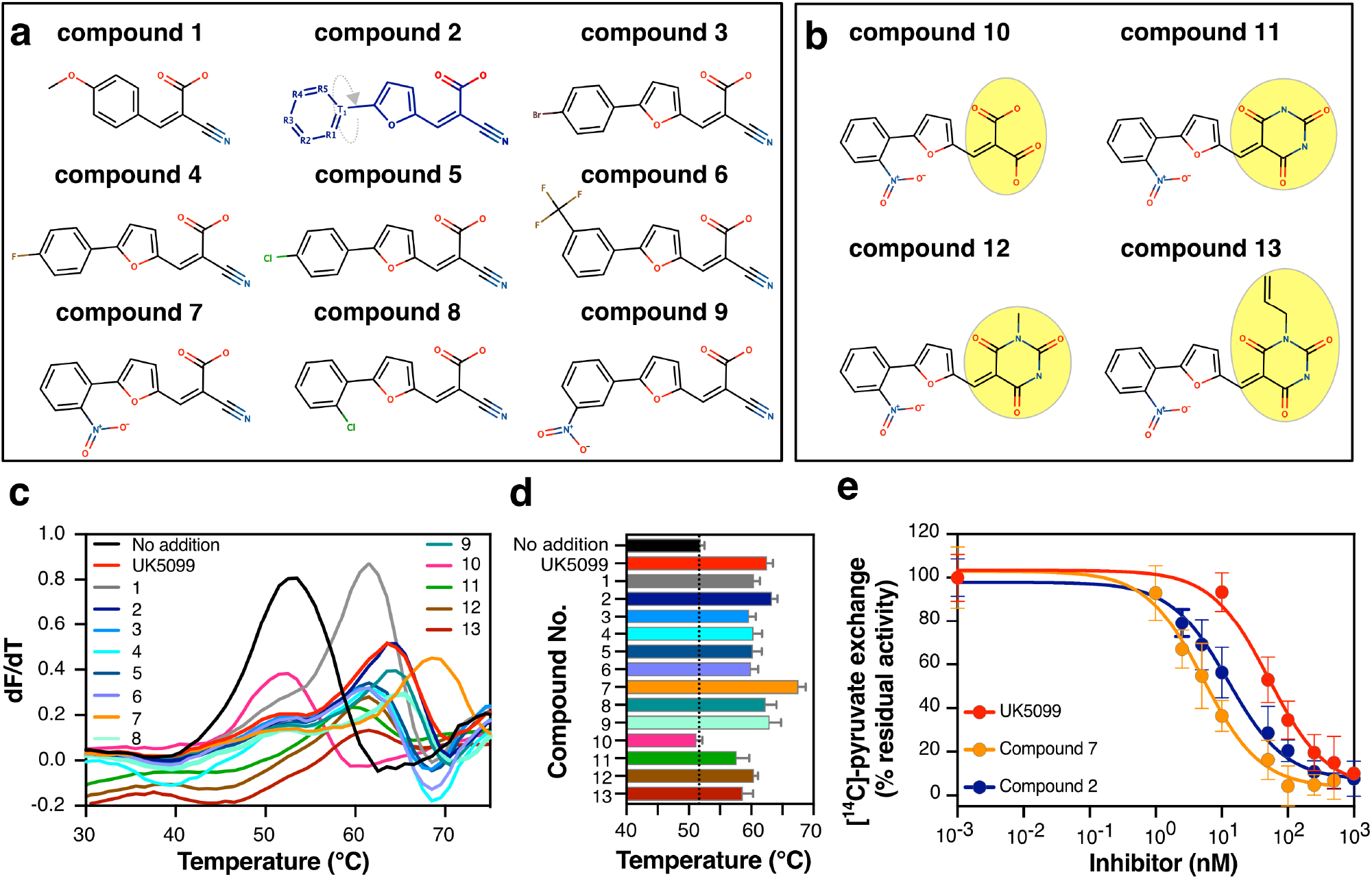
Chemical evolution of MPC inhibitors. (**a**) Chemical structures selected using the three hydrogen bond acceptors of UK5099. (**b**) Chemical structures of compounds where the cyanide and carboxylic acid groups have been replaced. (**c**) First derivatives obtained from CPM thermostability shift assays for the compounds in (**a**) and (**b**). (**d**) The Tm values, calculated from the first derivative in thermostability shift assays were plotted in a bar graph. The dotted line indicates the melting temperature of unliganded MPC. Color coding is as in (**c**). (**e**) Inhibition of [^14^C]-pyruvate homo-exchange by UK5099 (10–10,000 nM), compounds 2 (2.5–1,000 nM) and 7 (1–500 nM). Data information: In (**c**), data are representative of three biological repeats. Data in (**d**) have been calculated from three independent biological repeats (values included in **Table EV1**). In (**e**), data points represent the mean ± s.d of three biological repeats, each performed in triplicate. The averaged IC_50_-values were 52.6 ± 8.3 nM, 13.5 ± 4.1 nM and 5.4 ± 1.1nM for UK5099, compound 2 and 7, respectively.

To investigate the second aromatic ring further, we used the general chemical formula of compound 2 (**Figure 3a,** highlighted in blue) as a starting point and selected compounds with additional chemical groups on this aromatic ring. It has to be noted that R_1_ and R_2_ are equivalent to R_5_ and R_4_, respectively, due to a molecular torsion point (T_1_). Introduction of hydrophilic groups at the R_3_ position, as shown in compounds 3, 4, and 5 decreased binding compared to compound 2 and UK5099 (**Figure 3c and d** and **Table EV1**). However, in the case of compound 7, binding was enhanced by a nitro group in position R_1_ (or R_5_) (**Figure 3a**), which shifted the MPC thermostability to 67.8 ± 1.2 °C, more than the thermal shifts of compound 2 or UK5099 (Tm; compound 2: 63.4 ± 0.9 °C UK5099: 62.6 ± 0.9 °C) (**Figure 3c** and **d** and **Table EV1**). Compound 7 inhibited pyruvate transport in proteoliposomes, with an IC_50_-value of 5.4 ± 1.1 nM, approximately ten times lower than that of UK5099 (52.6 ± 8.3 nM) and two times lower than that of compound 2 (12.4 ± 4.6 nM) (**Figure 3e**), confirming an important role of the nitro group in position R_1_ (or R_5_) for binding.

To investigate whether the reactive nature of the nitro group in compound 7 contributes to its higher potency, we tested compound 8, in which a chloride group replaces the nitro group. This led to a thermostability shift lower than that of compound 7 (**Figure 3a, c-d** and **Table EV1**). Compound 9, in which the nitro group is relocated to position R_2_ (or R_4_), decreased the thermal shift compared to compound 7 but retained a thermal shift similar to that of compound 2 (**Figure 3a, c** and **d** and **Table EV1**). Replacement of the nitro group at position R_2_ (or R_4_) in compound 6 by a less reactive group reduced the thermal shift compared to compound 9 (**Figure 3b-d**). Therefore, the higher thermal shift elicited by compound 7 is possibly due to additional interactions introduced by the nitro group (Wilcken *et al*., 2013).

We evaluated the importance of the hydrogen bond acceptors provided by cyanide and carboxylic groups in compound 7 by testing compounds where these moieties are replaced by others (**Figure 3b**). First, we tested the replacement of the cyanide group by a carboxyl group (*i.e.,* compound 10). No thermal shift could be detected by compound 10 (**Figure 3c-d** and **Table EV1**), indicating that the interaction with MPC was abolished or very weak. This observation is in accordance with earlier observations showing a disruption of inhibitory capacity upon replacement of the cyanide group of UK5099 (Halestrap, 1975). This result could be explained by the introduction of a negatively charged and larger moiety at the position of the hydrogen bond acceptor (feature III, **Figure 3g**). None of the other inhibitors have a negative group in that position (**Figure EV4c-f** and **Figure 3a-b**). The inability of compound 10 to produce any thermostability shift despite the presence of all other chemical features of compound 7, including the activated double bond, emphasizes the importance of the hydrogen bond acceptors for binding to MPC.

Subsequently, we tested compounds that partly match the pyruvate-like pharmacophore. Both the cyanide and carboxyl group were replaced by a barbituric acid (compound 11), which matches with two of the three hydrogen bond acceptor features, by aligning with the ketone group (feature III) and carboxylic acid group (feature IV) of pyruvate (**Figure EV5**). We observed binding to MPC by this compound, albeit with smaller thermal shifts compared to UK5099, and by two closely related compounds (*i.e.,* compound 12 and 13), which have a slightly larger functional group (**Figure 3c-d** and **Table EV1**). This result demonstrates that matching two of the hydrogen acceptor features might be a minimal requirement for binding to MPC.

The above results highlight the importance of the three hydrogen bond acceptors and the adjacent aromatic rings as key chemical features for high potency, which could guide the identification of new MPC inhibitors. Most importantly, we present here new high affinity MPC inhibitors and we have identified compound 7 as the most potent one.

### High affinity inhibition is not attributed to covalent interactions with the human MPC

We then asked the question whether UK5099 and compound 7, which share inhibitory potencies in the low nanomolar range, form covalent interactions with MPC. A common characteristic of these compounds is an activated double bond, a putative Michael addition site. Previously, it has been hypothesized that UK5099 could inhibit MPC by using Michael addition to form a covalent bond with the thiol group of a cysteine (Halestrap, 1975, Halestrap, 1976, Yamashita *et al*., 2020). Cys54 in MPC2 has been proposed to engage with an irreversible MPC chemical probe (Yamashita *et al*., 2020), raising the question whether the high affinity inhibitors with activated double bonds also react covalently with this cysteine.

Two approaches were used to investigate the potential covalent interactions of MPC with UK5099 and derivatives. First, the molecular masses of unliganded human MPC1L/MPC2 were measured by electrospray mass spectrometry and compared with the same protein complex after inhibition with UK5099, compound 7, or zaprinast. The first two inhibitor compounds contain the activated double bond and zaprinast lacks it. Covalent modification of MPC by an inhibitor would increase the molecular mass of either MPC protomer. However, the mass spectra of all MPC complexes were identical in the presence or absence of inhibitors and no molecular mass differences in MPC protomers were observed (**Table 1**). However, when the thiol reactive compounds NEM or MTSEA were used as positive controls, both MPC subunits were modified covalently and mass differences associated with modification of cysteine residues were observed (**Table 1**). These results demonstrate clearly that the inhibitor interactions with MPC are not covalent.

**Table 1:**
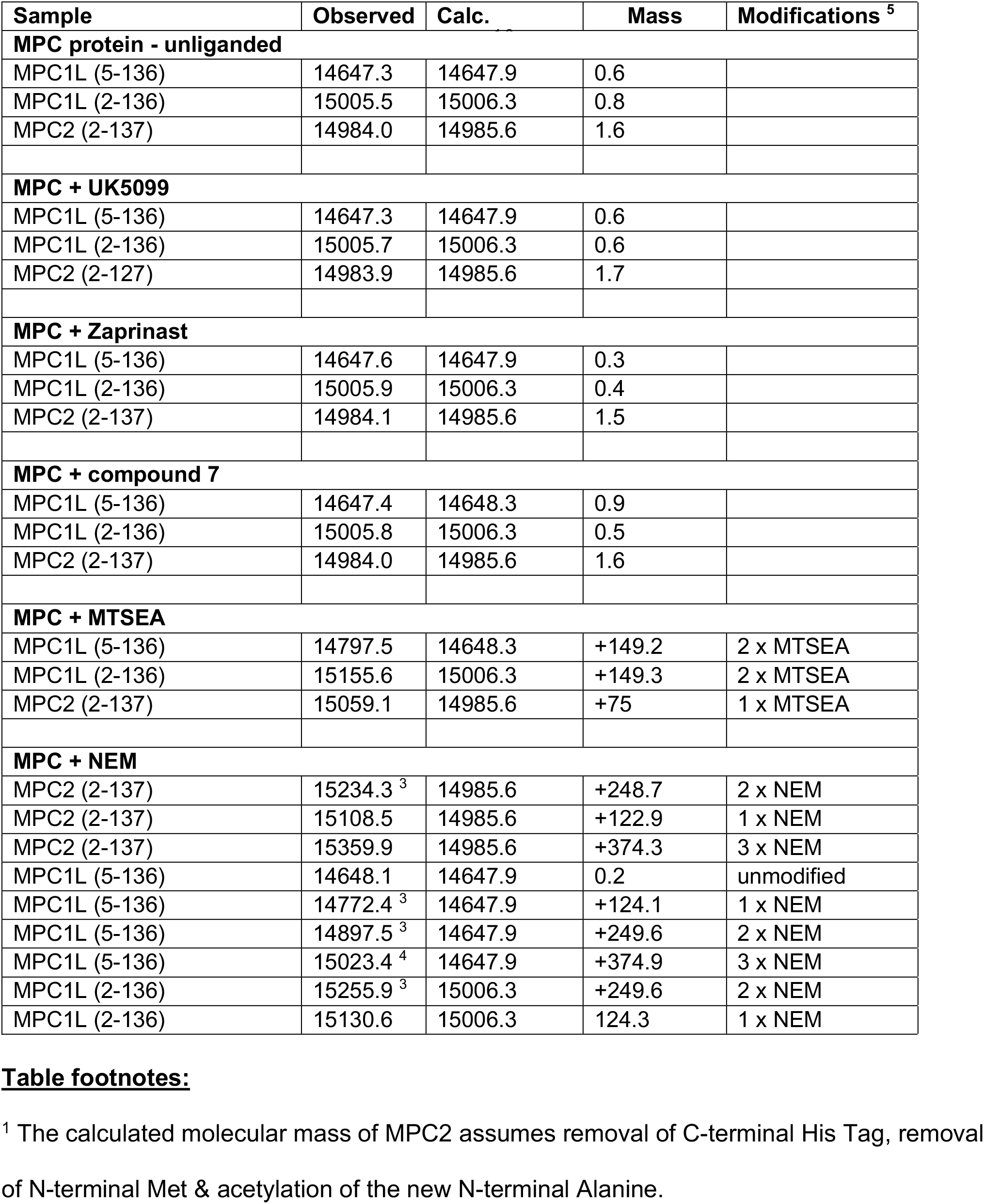
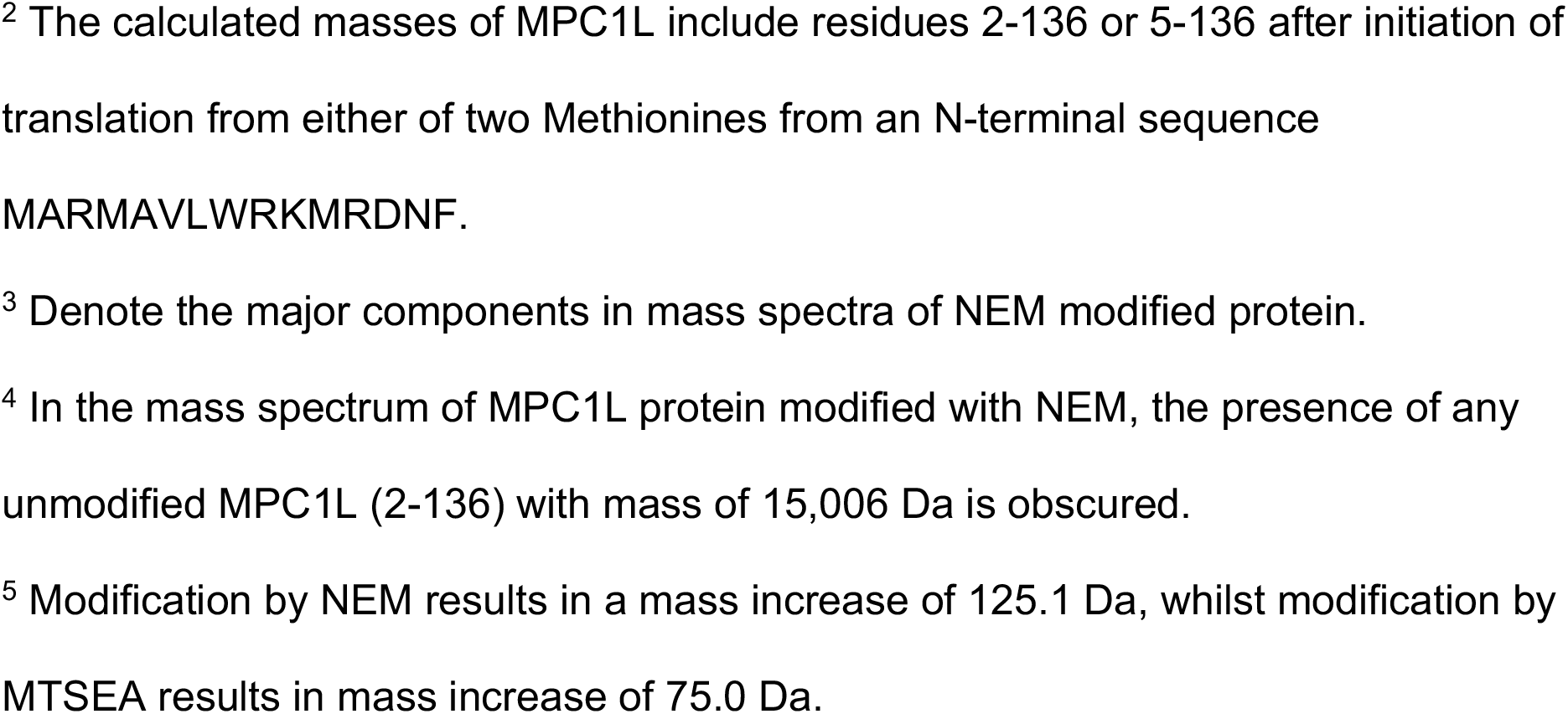
Molecular mass measurements of components of recombinant human MPC1L/MPC2 after reaction with either inhibitors or sulfhydryl modifiers

| Sample | Observed | Calc. | Mass | Modifications <sup>5</sup> |
| --- | --- | --- | --- | --- |
| **MPC protein - unliganded** |  |  |  |  |
| MPC1L (5-136) | 14647.3 | 14647.9 | 0.6 |  |
| MPC1L (2-136) | 15005.5 | 15006.3 | 0.8 |  |
| MPC2 (2-137) | 14984.0 | 14985.6 | 1.6 |  |
| **MPC + UK5099** |  |  |  |  |
| MPC1L (5-136) | 14647.3 | 14647.9 | 0.6 |  |
| MPC1L (2-136) | 15005.7 | 15006.3 | 0.6 |  |
| MPC2 (2-127) | 14983.9 | 14985.6 | 1.7 |  |
| **MPC + Zaprinast** |  |  |  |  |
| MPC1L (5-136) | 14647.6 | 14647.9 | 0.3 |  |
| MPC1L (2-136) | 15005.9 | 15006.3 | 0.4 |  |
| MPC2 (2-137) | 14984.1 | 14985.6 | 1.5 |  |
| **MPC + compound 7** |  |  |  |  |
| MPC1L (5-136) | 14647.4 | 14648.3 | 0.9 |  |
| MPC1L (2-136) | 15005.8 | 15006.3 | 0.5 |  |
| MPC2 (2-137) | 14984.0 | 14985.6 | 1.6 |  |
| **MPC + MTSEA** |  |  |  |  |
| MPC1L (5-136) | 14797.5 | 14648.3 | +149.2 | 2 x MTSEA |
| MPC1L (2-136) | 15155.6 | 15006.3 | +149.3 | 2 x MTSEA |
| MPC2 (2-137) | 15059.1 | 14985.6 | +75 | 1 x MTSEA |
| **MPC + NEM** |  |  |  |  |
| MPC2 (2-137) | 15234.3 <sup>3</sup> | 14985.6 | +248.7 | 2 x NEM |
| MPC2 (2-137) | 15108.5 | 14985.6 | +122.9 | 1 x NEM |
| MPC2 (2-137) | 15359.9 | 14985.6 | +374.3 | 3 x NEM |
| MPC1L (5-136) | 14648.1 | 14647.9 | 0.2 | unmodified |
| MPC1L (5-136) | 14772.4 <sup>3</sup> | 14647.9 | +124.1 | 1 x NEM |
| MPC1L (5-136) | 14897.5 <sup>3</sup> | 14647.9 | +249.6 | 2 x NEM |
| MPC1L (5-136) | 15023.4 <sup>4</sup> | 14647.9 | +374.9 | 3 x NEM |
| MPC1L (2-136) | 15255.9 <sup>3</sup> | 15006.3 | +249.6 | 2 x NEM |
| MPC1L (2-136) | 15130.6 | 15006.3 | 124.3 | 1 x NEM |

Secondly, we mutated each cysteine of MPC2 (Cys54) or MPC1L (Cys85 or Cys87) separately to alanine. **Figure 4a-b** shows the purification and unfolding profiles of the three cysteine-to-alanine mutants, demonstrating that each one of them was folded in detergent/lipid-containing buffer. We reasoned that if the low nanomolar potencies of UK5099 or compound 7 are due to covalent bond formation, then abolishing such a critical interaction by mutation of an interacting cysteine to alanine should translate to a broad decrease in binding and therefore, a decrease in the observed thermostability shifts in the presence of inhibitor. We compared wild type and cysteine-to-alanine mutants for the thermostability shift elicited by UK5099, CHC and compound 7, representing compounds with an activated double bond (**Figure 4c**). We also tested zaprinast and the TZD mitoglitazone, representing compounds without the activated double bond. For each of these mutants, we observed thermostability shifts comparable to wild type with all tested compounds (**Figure 4c**). Interestingly, mutating Cys54 to alanine in MPC2 did not reduce, but rather increased the thermostability shifts, which is not compatible with this residue participating in any critical interaction with these inhibitors. This result indicates an indirect, positive effect on binding by reducing the size of the side chain. The thermostability shift was also not abolished when each of the cysteines in MPC1L were mutated to alanine. Taken together, these results challenge previous proposals for cysteine-mediated covalent bonds between MPC and specific inhibitors.

**Figure 4.**
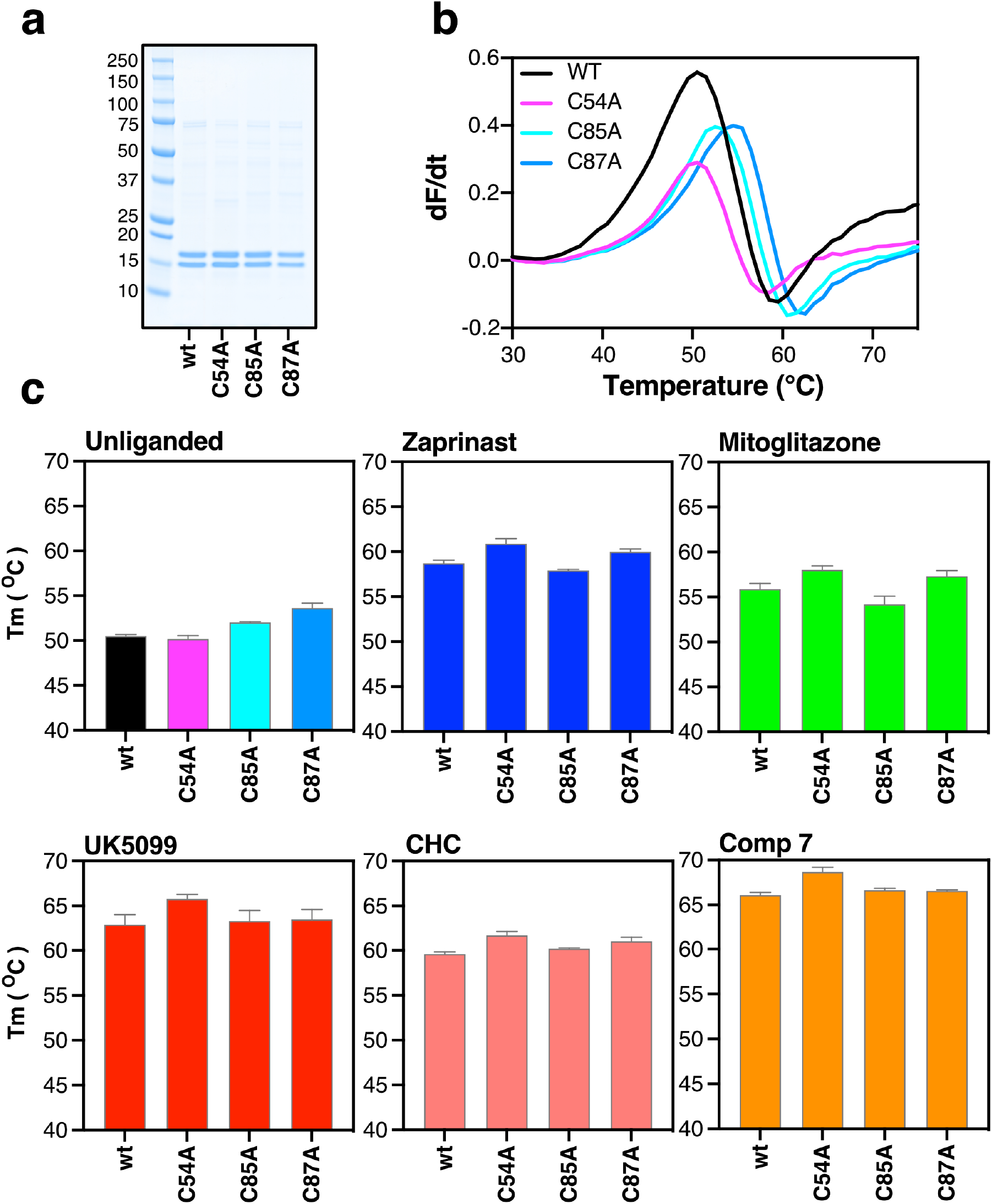
MPC cysteines do not form covalent interactions with inhibitors. (**a**) Cysteine-to-alanine replacement mutants purified in detergent in parallel with wild-type (wt) human MPC1L/MPC2 via nickel affinity chromatography. (**b**) First derivative of protein unfolding profiles for wild-type and cysteine-to-alanine replacement mutants via CPM thermostability assay. (**c**) CPM thermostability for wild-type and cysteine-to-alanine replacement mutants in the absence or presence of zaprinast, and mitoglitazone, which do not feature the activated double bond, or CHC, UK5099 and compound 7, each containing an activated double bond. Data information: In (**b**) data are representative of two biological repeats and three technical repeats. In (**c**), data represent the mean ± s.d of two biological repeats and three technical repeats.

### An extended pharmacophore model successfully identifies commonly used drugs as potent MPC inhibitors

Having established the essential features for MPC inhibition, we tested whether their presence could be used to identify new compounds that inhibit MPC. We created a refined pharmacophore model where we incorporated all features resulting in the increased affinity of compound 7 (**Figure 5c** and **g**), including an additional aromatic structure (feature VIII) and two hydrogen bond acceptors (features IX and X). Importantly, both aromatic rings of UK5099 overlap with the ones in compound 7, which may be contributing to the high potency of these inhibitors (**Figure 5c** and **g** and **Figure EV6**).

**Figure 5.**
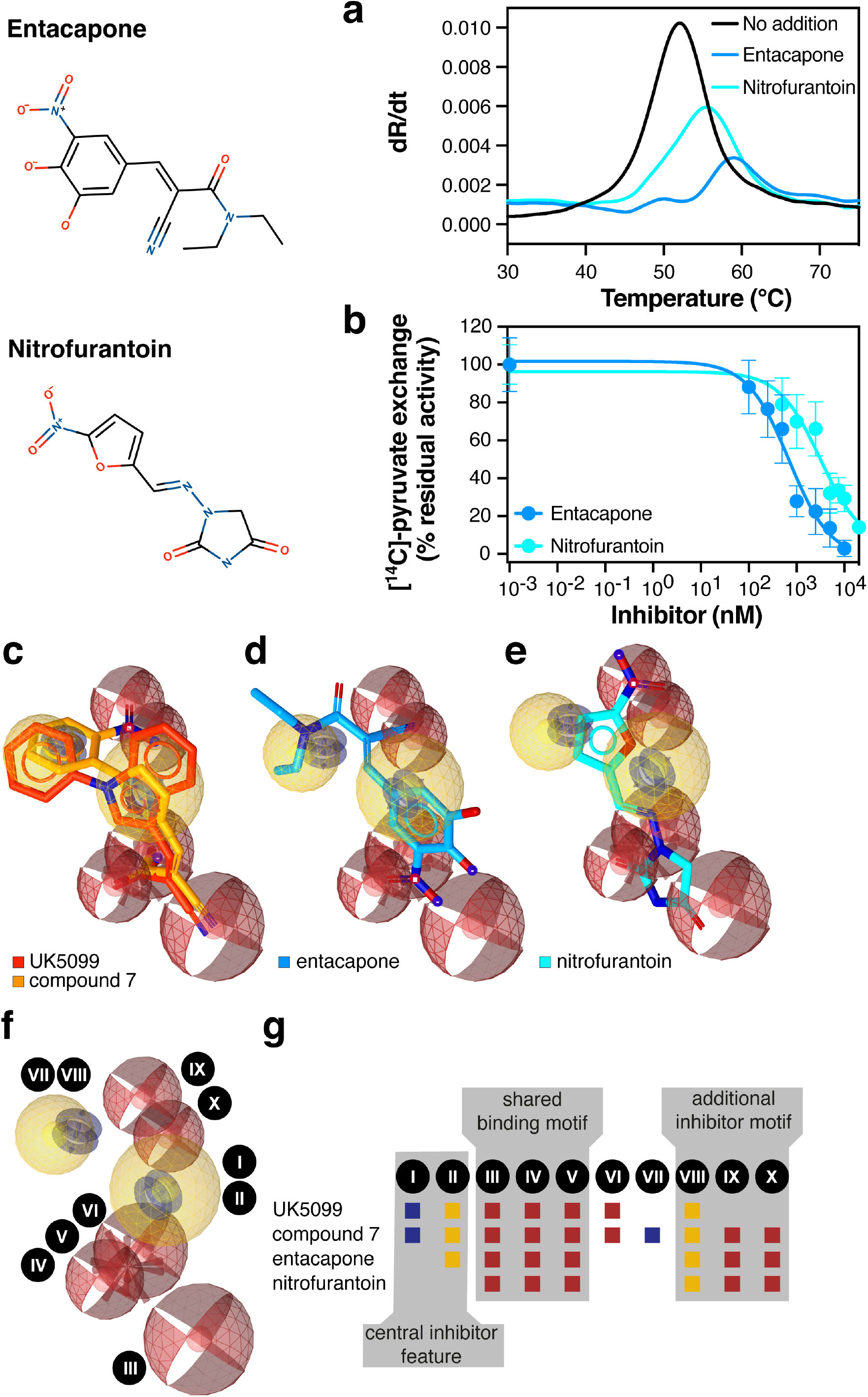
Matching of an extended pharmacophore model predicts MPC inhibition. (**a**) Thermostability assays for nitrofurantoin and entacapone using nano-DSF. (**b**) Inhibition of [^14^C]-pyruvate homo-exchange by entacapone (100–10,000 nM) and nitrofurantoin (500–20,000 nM). (**c**) Extended pharmacophore features of compound 7 and comparison with UK5099. Red spheres indicate hydrogen acceptor features, yellow spheres hydrophobic ring features, red star indicate negative ionizable features, and bleu tori indicate aromatic ring features. (**d**) Fitting of nitrofurantoin in the extended pharmacophore. (**e**) Fitting of entacapone in the extended pharmacophore. (**g**) 3D representation of the extended pharmacophore with the identified features (roman numbers) and their presence in the various compounds. Data information: In (**a-b**), data are representative of two biological repeats, each performed in triplicate.

We asked whether this model can be used to identify off-target effects of existing small molecule drugs on MPC. Several medicines in development or in clinical practice suffer from mitochondrial toxicity but their mitochondrial targets are usually unknown. In this way, we have identified two commonly used compounds as potent MPC inhibitors, reported previously to interfere with mitochondrial function (Carbonera *et al*., 1988, Grunig *et al*., 2017, Grunig *et al*., 2018). The first one, entacapone ((2*E*)-2-Cyano-3-(3,4-dihydroxy-5-nitrophenyl)-*N*,*N*-diethylprop-2-enamide), is used in combination therapy for Parkinson’s disease and the other, nitrofurantoin ((*E*)-1-[(5-nitro-2-furyl)methylideneamino]imidazolidine-2,4-dione), is an antibiotic used to treat bladder infections. Interestingly, the pharmacophore features of both drugs fit the essential pyruvate-like binding motif as well as the additional binding motif, similar to compound 7 (**Figure 5d-g** and **Figure EV6**).

Both compounds were tested on purified MPC1L/MPC2 using nano-DSF (**Figure 5a**), as they both had quenching effects on CPM fluorescence. Interestingly, entacapone, which contains all the essential features of the extended pharmacophore, shifted the MPC unfolding curve from 51.9 ± 0.1 °C to 58.2 ± 1.0 °C whilst a more moderate shift to 54.7 ± 0.2 °C was observed with nitrofurantoin (**Figure 5a** and **Table EV1**). Consistent with these results, the inhibitory potency of entacapone in MPC1L/MPC2 proteoliposomes was 630 ± 53 nM, whereas the IC_50_ for nitrofurantoin was in the low micromolar range (3.3 ± 2.6 μM). The decreased potency of nitrofurantoin compared to entacapone could be explained by the absence of the central aromatic ring moiety (**Figure 5f** and **g**). Therefore, the proposed key chemical features for MPC inhibition can guide the identification of new or previously unidentified small molecule inhibitors.

## DISCUSSION

The molecular identification of MPC has sparked an interest to explore the potential of this key metabolic protein as a drug target but, in the absence of structural information, our understanding of pyruvate transport and inhibition remains limited. Different claims have been made regarding the ability of MPC homo- and hetero-complexes to support transport and binding (Compan *et al*., 2015, Lee *et al*., 2020, Nagampalli *et al*., 2018, Tavoulari *et al*., 2019, Vanderperre *et al*., 2016). We have demonstrated the formation of the two proposed human hetero-dimers by co-expressing and co-purifying tagged MPC2 with untagged MPC1 or MPC1L and we have reconstituted the human hetero-dimer MPC1L/MPC2 into functional liposomes, providing evidence that hetero-dimer formation mediates robust pyruvate transport activity. This result is consistent with our previous study of the yeast MPC complexes (Tavoulari *et al*., 2019) and studies by others showing that expression of both MPC proteins is necessary for pyruvate transport in mitochondria (Bricker *et al*., 2012, Compan *et al*., 2015, Herzig *et al*., 2012).

We showed that both hetero-dimers bind known MPC inhibitors, whereas MPC1 alone could not be purified and MPC2 did not bind small molecules on its own. This notion is in contrast to previous reports claiming binding of inhibitors to MPC2 homomers (Lee *et al*., 2020, Nagampalli *et al*., 2018). Notably, the binding affinities of known inhibitors to MPC1 or MPC2 homomers, proposed by others, were in the micromolar range (Lee *et al*., 2020), whereas studies in isolated mitochondria had predicted affinities in the low nanomolar range for mammalian MPCs (*i.e.,* K_i_ for UK5099 was ∼50 nM in isolated mitochondria (Halestrap, 1975)). Consistent with the studies in mitochondria, we report inhibitory potencies in the low nanomolar range for the hetero-dimer (*i.e.,* IC_50_-value for UK5099 is 52.6 ± 8.3 nM in our study). It is most likely that maximal binding occurs in the interface between protomers but it cannot be excluded that MPC1 contributes essential coordinating residues.

An interesting observation is that the MPC1L/MPC2 hetero-dimer, expressed in testis, binds and is inhibited by compounds (**Figure 2**) previously identified for their ability to inhibit the ubiquitously expressed MPC1/MPC2 hetero-dimer in human cells. This observation supports the idea that the two hetero-dimers should share a similar binding site, in agreement with their high sequence conservation (**Figure EV1**). On the contrary, the inhibitory potencies of the tested compounds were 25 to 200 times higher for the human MPC1L/MPC2 compared to the yeast Mpc1/Mpc3, suggesting important differences in inhibitor co-ordination between species.

From the inhibitors tested in our system, UK5099 was clearly the one with the highest potency of all previously reported compounds. The second most potent inhibitor was zaprinast, followed by the anticancer agent lonidamine and the insulin sensitizer mitoglitazone. Consistent with our work, lonidamine inhibits MPC in isolated mitochondria with a K_i_ at 2.5 μM (Nancolas *et al*., 2016). The high affinity of *α*-cyano-cinnamates and derivatives for MPC has been previously attributed to an activated double bond, which could potentially form Michael addition adducts with cysteine residues in MPC. Cys54 in MPC2 can engage with an irreversible chemical probe, which is chemically different than UK5099 (Yamashita *et al*., 2020) but this raised the possibility that Cys54 also engages covalently with UK5099. A covalent interaction could also explain the high potencies of UK5099 and derivatives relative to other MPC inhibitors. However, we show here that the nanomolar potency of these compounds is not attributed to the formation of covalent bonds. No cysteine modifications were observed by ESI-MS molecular mass measurements of MPC after inhibition. Replacement of Cys54 of MPC2 to alanine did not prevent, but rather increased binding of different inhibitors tested. Replacement of each of the cysteines in MPC1L also resulted in similar thermostability compared to wild-type in the presence of any inhibitor.

Understanding the common pharmacophore properties of MPC inhibitors allowed us to identify more MPC binders and improve the inhibitory potency. Our pharmacophore analysis has revealed that substrate and small molecule inhibitors share an arrangement of three hydrogen bond acceptors. In the substrate, the hydrogen bond acceptors are provided by carboxyl and ketone groups. In agreement, MPC has also been implicated in the transport of the ketone bodies acetoacetate and β-hydroxybutyric acid (Papa & Paradies, 1974, Paradies & Papa, 1975). In inhibitors, the three hydrogen bond acceptors can be supplied by a carboxyl group and a nitrile group, as in UK5099 and compounds 1 to 9. Consistent with the importance of the carboxyl and nitrile groups as efficient hydrogen bond acceptors, *α*-fluorocinnamate is 1,000 times less potent than *α*-cyano-cinnamate (Halestrap, 1975).

Distinct between inhibitors and substrates, but common amongst all inhibitors, is a central aromatic ring moiety that appears to be the second requirement for MPC inhibition, after the hydrogen bond acceptors. A second aromatic ring moiety is an additional feature that might be adding to the number of interactions between inhibitor and MPC to increase affinity. Moreover, addition of a nitro group to this second moiety led to a molecule with increased inhibitory potency (compound 7). These new features allowed us to understand binding of two commonly used drugs, entacapone and nitrofurantoin and could guide the identification of additional off-target effects on MPC.

We have described the development of *in vitro* assays for MPC and have shown the use of a thermostability shift assay as a primary screening high-throughput approach allowing high affinity MPC binders to be identified. In combination with a proteoliposome-based transport assay, thermostability shift assays have allowed us to identify new small molecule inhibitors and extend the potency of previously known ones to lower nanomolar range. By combining *in vitro* and *in silico* analyses we have mapped the pharmacophore properties of MPC binders and propose that a set of three hydrogen bond acceptors followed by a hydrophobic moiety are minimal requirements for high affinity binding, whereas covalent bonds are not formed between MPC and its inhibitors. This knowledge could be key to accelerate efforts for the development of clinically relevant MPC modulators.

## MATERIALS AND METHODS

### Compounds

UK5099, CHC, zaprinast, lonidamine, mitoglitazone, pioglitazone, rosiglitazone, valproic acid, silibinin, entacapone and nitrofurantoin were purchased from Sigma Aldrich (Merck, Gillingham, UK). UK5099-based novel compounds (*i.e.,* compounds 1-13) were obtained from the Chembridge chemical store Hit2Lead (ChemBridge Corporation, CA, USA). All compounds were dissolved at a 100 mM stock concentration in DMSO.

### Molecular biology

The codon-optimized gene sequences for human MPC1 (UniProt: Q9Y5U8), MPC2 (UniProt:O95563) and MPC1L (UniProt: P0DKB6) were synthesized (GenScript, Piscataway, NJ, USA) and cloned into the bidirectional expression vector pBEVY-GU (gift from Charles Miller; Addgene plasmid # 51229 (Miller *et al*., 1998)). For expression of *mpc1* or *mpc1L*, the cDNA was subcloned into the EcoRI/SacI sites and for *mpc2* into the BamHI/XbaI sites. The sequence of MPC2 was designed to include a sequence coding for Factor Xa or TEV protease cleavage sites, followed by a tag of eight-histidines at their C-termini.

### Protein expression and mitochondrial preparations

The expression plasmids were transformed into the *mpc* triple deletion strain of *S. cerevisiae* SHY15 (Herzig *et al*., 2012), kindly provided by Jean-Claude Martinou, using standard methods (Gietz & Schiestl, 2007). Successful transformants were selected on synthetic-complete uracil-dropout medium plates supplemented with 2% (*w/v*) glucose. Pre-cultures were grown in the same medium and used to inoculate 50 L of YPG medium containing 0.1% (*w/v*) glucose in an Applikon Pilot Plant 140 L bioreactor (Thangaratnarajah *et al*., 2014). Protein expression was induced for 3 h with 0.4% (*w/v*) galactose, after 20 h of growth in YPG. Mitochondrial isolation was performed, as previously described (Thangaratnarajah *et al*., 2014), using a DYNO-MILL (Willy A. Bachofen AG, Basel, Switzerland). Isolated crude mitochondria were aliquoted, flash-frozen in liquid nitrogen and stored in −80 °C until use.

### Affinity chromatography

The MPC1L/MPC2 and MPC1/MPC2 hetero-complexes were purified from mitochondria of the SHY15 strain expressing the relevant protein pairs. Immediately prior to purification, 1 g of mitochondria were thawed and suspended in buffer containing 20 mM Tris-HCl, pH 7.4, 150 mM NaCl, 10% (*v/v*) glycerol, one Complete EDTA-free protease inhibitor cocktail tablet (Roche, Basel, Switzerland) and 1% (*w/v*) lauryl maltose neopentyl glycol (LMNG, Anatrace, OH, USA). Mitochondria were solubilized for 1.5 h at 4 °C under gentle agitation and the insoluble material was removed by ultracentrifugation at 205,000 x *g* for 45 min. The supernatant was incubated for 2 h with nickel Sepharose High Performance beads (Cytiva, MA, USA), in the presence of 10 mM imidazole, and then transferred into an empty column (Bio-Rad, Watford, UK). The column was washed initially with 25 column volumes of buffer A (20 mM Tris-HCl, pH 7.4, 150 mM NaCl, 50 mM imidazole, 0.1% (*w/v*) LMNG, 0.1 mg/mL tetraoleoyl cardiolipin (TOCL, Avanti Polar Lipids, AL, USA)), followed by 20 column volumes of buffer B (20 mM Tris-HCl, pH 7.4, 150 mM NaCl, 0.1% (*w/v*) LMNG, 0.1 mg/mL TOCL). MPC1/MPC2 was eluted from the column by on-column digestion, in the presence of 5 mM CaCl_2_, for 12 h at 4 °C with 10 μg of Factor Xa protease (New England Biolabs, Hitchin, UK) per 1 g of mitochondria. MPC1L/MPC2 was eluted by on-column digestion for 12 h at 10 °C with 250 μg of MBP-TEV protease per 1 g of mitochondria. The mobile phase containing untagged MPC was separated from the resin with empty Proteus Midi spin columns (Generon, Slough, UK) at 200 x *g* for 5 min. For MPC1L/MPC2, the TEV protease was removed by incubation of the purified protein with 250 µL amylose resin (New England Biolabs, Hitchin, UK). Protein concentrations were determined by the bicinchoninic acid assay (Thermo Fisher Scientific, Hemel, Hempstead, UK).

### Size-exclusion chromatography

Analytical size-exclusion chromatography was performed on an ÄKTA Explorer (GE Healthcare) with a Superdex 200 10/300 GL column (GE Healthcare) equilibrated in SEC buffer (20 mM Tris-HCl, pH 7.4, 150 mM NaCl, 0.05% (w/v) LMNG, 0.05 mg/ml TOCL). Nickel purified MPC1L/MPC2 or Mpc1/Mpc3 were applied onto the column at 0.3 ml/min and 0.15 ml fractions were collected. The column was calibrated with a high molecular weight calibration kit (GE Healthcare) in the same buffer without detergent and lipid.

### Mass determination by SEC-MALLS

SEC-MALLS analysis was performed with a Superdex 200 10/300 GL column (GE Healthcare) on an ÄKTA Explorer (GE Healthcare) coupled in-line with a light scattering detector (Dawn HELEOSII, Wyatt Technologies) and a refractometer (Optilab T-rREX, Wyatt Technologies). The MPC complexes were applied onto the Superdex 200 10/300 GL column equilibrated with 20 mM Tris-HCl, pH 7.4, 150 mM NaCl, 0.005% (w/v) LMNG, 0.005 mg/ml TOCL at 0.3 ml/min. All data were recorded and analysed with ASTRA 6.03 (Wyatt Technologies). Molecular weight calculations were performed using the protein-conjugate method (Slotboom *et al*., 2008) with the dn/dc value for protein of 0.185 ml/g and dn/dc value for LMNG-TOCL of 0.1675 ml/g (Thangaratnarajah *et al*., 2014). The contributions of each protein to the overall protein-detergent-lipid complex were determined from the extinction coefficients *ε*_A280_, derived from the amino-acid sequence using the ProtParam tool on the ExPaSy server (Gasteiger *et al*., 2005).

### Thermostability shift analysis

The assessment of ligand binding was performed via shifts in protein thermostability upon binding using thermal denaturation on a rotary quantitative PCR (qPCR) instrument (Qiagen Rotor-Gene Q 2plex HRM, Venlo, Netherlands), as previously described (Crichton *et al*., 2015). In this method, cysteine residues, buried within the protein structure, become solvent exposed during denaturation in a temperature ramp and react with 7-diethylamino-3-(4-maleimidophenyl)-4-methylcoumarin (CPM) to form fluorescent-adducts. Briefly, a CPM working solution was prepared by diluting the CPM stock (5 mg/ml in dimethyl sulfoxide) 50-fold into assay buffer (20 mM Tris-HCl, pH 7.4, 150 mM NaCl, 0.1% (*w/v*) LMNG, 0.1 mg/ml TOCL) and incubated for 10 min at room temperature. In each analysis, performed in duplicates or triplicates, 3 μg of purified MPC was diluted into 45 μL of assay buffer containing the desired concentration of the inhibitor, incubated for 10 min on ice and then 5 μL of the CPM working solution were added. Samples were incubated on ice for a further 10 min and then subjected to a temperature ramp of 5.6 °C/min. The fluorescence increase was monitored with the HRM channel of the instrument (excitation at 440–480 nm, emission at 505–515 nm). Unfolding profiles were analyzed with the Rotor-Gene Q software 2.3 and the peaks of their derivatives were used to determine the apparent melting temperature.

Ligand binding was also assessed using dye-free nano differential scanning fluorimetry (nano-DSF), which monitors fluorescence changes due to altered environments of tryptophan and tyrosine residues during unfolding. Protein samples in buffer containing 20 mM Tris-HCl pH 7.4, 150 mM NaCl, 0.1% (*w/v*) LMNG, 0.1 mg/mL TOCL, in the presence or absence of the indicated concentrations of small molecule inhibitors, were loaded into capillary tubes, and a temperature ramp of 5 °C/min was applied. The intrinsic fluorescence was measured using the NanoTemper Prometheus NT.48 (NanoTemper, München, Germany).

### Reconstitution in proteoliposomes

Egg L-α-phosphatidylcholine 99% (*w/v*) (Avanti Polar Lipids, AL, USA) and TOCL were mixed in a 20:1 (*w/w*) ratio, dried under a stream of nitrogen and washed once with methanol before being dried again. Lipids were hydrated in 20 mM Tris-HCl, pH 8.0, 50 mM NaCl to a concentration of 12 mg/mL. Unlabeled pyruvate was added as a freshly made concentrated stock at a final concentration of 5 mM for internalization. Lipids were solubilized with 1.2% (*v/v*) pentaethylene glycol monodecyl ether (Sigma) and freshly purified protein was added at a lipid-to-protein ratio of 250:1 (*w/w*). Samples were incubated on ice for 5 min, and then liposomes were formed by the step-wise removal of pentaethylene glycol monodecyl ether by five additions at 20 min intervals of 60 mg Bio-Beads SM-2 (Bio-Rad, Watford, UK) with gentle mixing at 4 °C. A final addition of 480 mg Bio-Beads was incubated with the samples overnight. Proteoliposomes were first separated from the Bio-Beads by passage through empty spin columns (Bio-Rad, Watford, UK) and subsequently pelleted at 120,000 x *g* for 60 min. The proteoliposomes were resuspended with a thin needle in 150 μL of their supernatant after the rest of the supernatant was removed.

### Pyruvate transport assays

The time course of pyruvate homo-exchange was measured at room temperature. The transport was initiated by diluting proteoliposomes 200-fold into external buffer, 20 mM MES, pH 6.4, 50 mM NaCl, containing 50 μM [^14^C]-pyruvate (500,000 dpm, Perkin Elmer, MA, USA). The reaction (0–30 s) was terminated by rapid dilution into 8 volumes of ice-cold internal buffer (20 mM Tris-HCl, pH 8.0, 50 mM NaCl), followed by rapid filtration through cellulose nitrate 0.45 μm filters (Millipore, Gillingham, UK) and washing with an additional 8 volumes of buffer. The filters were dissolved in Ultima Gold scintillation liquid (Perkin Elmer, MA, USA) and the radioactivity was counted with a Perkin Elmer Tri-Carb 2800 RT liquid scintillation counter. For inhibition of pyruvate transport, various concentrations of compounds, as indicated in the figure legends, were added to the liposomes simultaneously with 50 μM radioactive substrate. The data analysis was performed with non-linear regression fittings using GraphPad Prism 7.0d ([Inhibitor] vs response, variable slope). The specific uptake rates were calculated based on the amount of protein used in reconstitutions, as estimated from bicinchoninic acid assay. The biological repeats represent independent proteoliposome preparations using protein from independent purifications.

### Pharmacophore modeling

All molecules were prepared for pharmacophore modelling, by setting the protonation at pH 7.4, which was also used for the thermostability assays, using Marvin Sketch (ChemAxon, Budapest, Hungary). Subsequently, pharmacophore models were generated using Ligandscout 4.3 Essential (Inte:ligand, Vienna, Austria). Three-dimensional structures were generated initially and energy was minimized using the MMFF94 protocol (see **Supplementary Table EV2** for minimized energies). Secondly, conformations were generated using the iCon protocol, allowing a maximum of 200 configurations per molecule with an energy window of 20 kcal/mol and a root mean square (RMS) value of 0.8 (see **Supplementary Table EV2** for number of conformations generated). Next, maximally ten shared-feature pharmacophore models were generated for each individual molecule using the ‘pharmacophore fit’ scoring function. The coordinates of the pharmacophores were determined and used to calculate distances between features. To investigate pharmacophore features shared between different compounds, pharmacophore models were transferred to the screening interface. A screening library containing all compounds was constructed using the same conformation generation settings, as described above, and a fit-score was calculated using the ‘pharmacophore fit’ scoring function.

### Mass spectrometry

Different MPC inhibitors (**Table1**) or sulfhydryl reagents were first incubated with MPC1L/MPC2 in a folded state and then the protein molecular masses were measured by electrospray mass spectrometry. The MPC subunits (ca 10 µg) were resolved by reverse-phase HPLC on a PLRP-S column (1.0 mm i.d. x 75 mm; VARIAN) using a gradient of 2-propanol in a solvent system compatible with the recovery of membrane proteins (Carroll et al., 2009). The eluate (50 µL/min) was introduced into a QTRAP4000 (SCIEX) mass spectrometer operating in positive ion single MS mode. The instrument was scanned from 700 m/z to 2300 m/z after calibration with ions from a mixture of 1.0 µM of myoglobin and 1.5 µM trypsinogen. Molecular masses were obtained by reconstruction of the spectra of multiply charged ions using PeakView (SCIEX).

### Data availability

Source data have been uploaded to the Dryad repository (doi:10.5061/dryad.stqjq2c2x).

## SUPPLEMENTARY INFORMATION

**Figure EV1.**
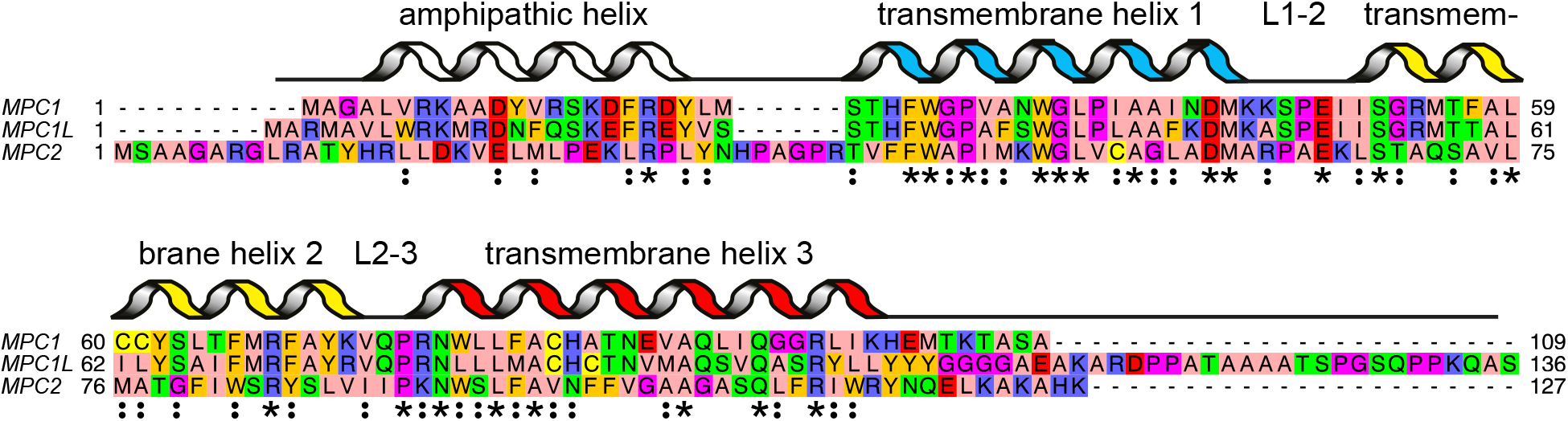
Sequence alignment and secondary structure prediction of human MPC proteins. The alignment was generated by Clustal Omega (Sievers *et al*., 2011), followed by manual curation. The aligned residues are colored by the ZAPPO color scheme in which aliphatic, polar, aromatic, positively charged, negatively charged, Pro/Gly and Cys, are colored pink, green, orange, blue, red, magenta and yellow, respectively. The asterisks indicate identical residues and the colon conserved substitutions. Also indicated are putative transmembrane helices, loop regions and the N-terminal amphipathic helix. The secondary structure elements were assigned based on PSIPRED (Buchan *et al*., 2013), MEMSAT3 (Jones *et al*., 1994) and conservation analysis.

**Figure EV2.**
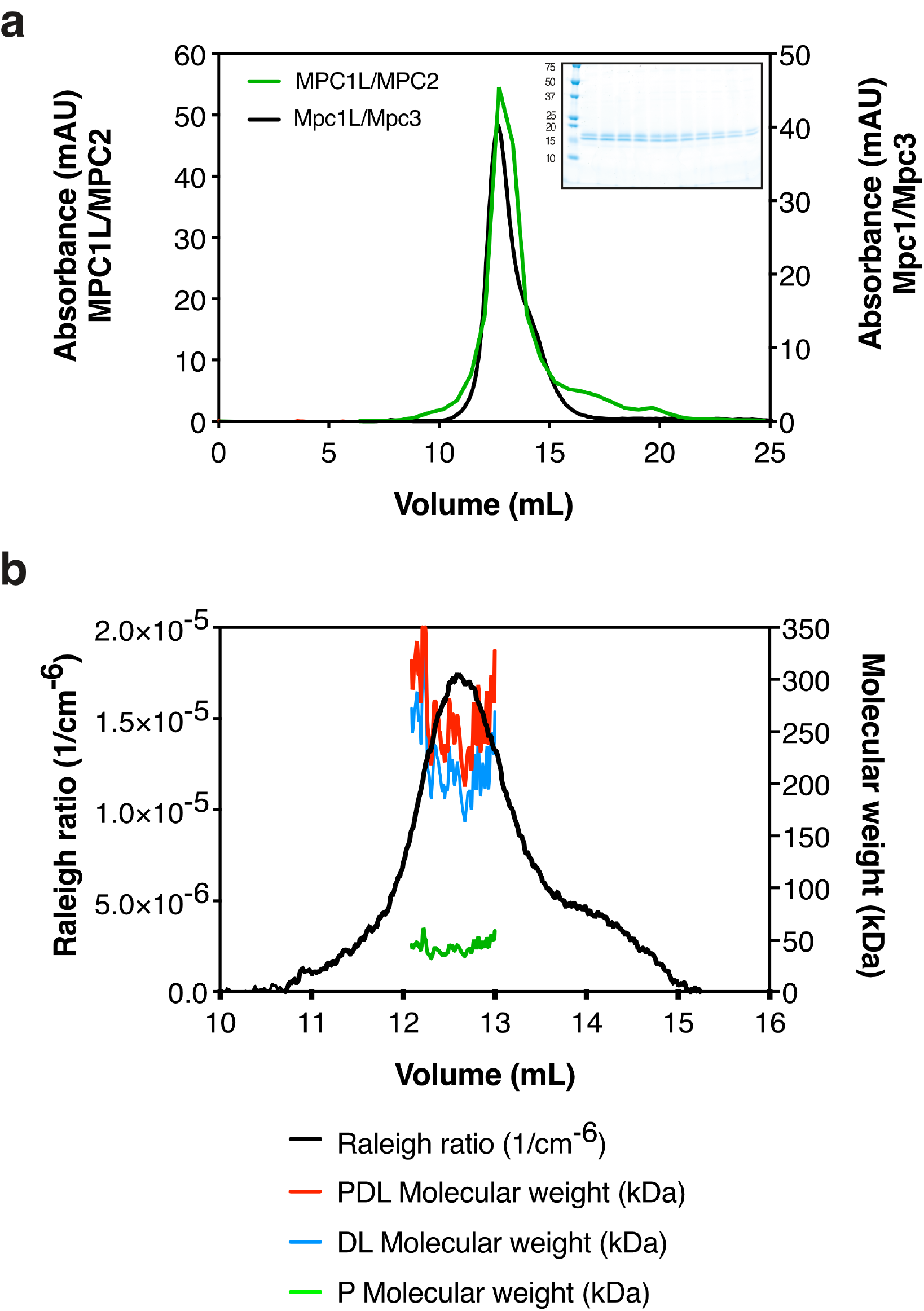
Oligomeric state of human MPC1L/MPC2. (**a**) Size-exclusion chromatography and superimposed A_280_ profiles for the Mpc1/Mpc3 hetero-dimer (Tavoulari *et al*., 2019) (black) and MPC1L/MPC2 (green) purified via Nickel-affinity chromatography. *Inset:* Peak fractions for MPC1L/MPC2 analyzed by SDS-PAGE and visualized by Coomassie Blue stain. (**b**) SEC-MALLS analysis of MPC1L/MPC2. The light scattering trace is shown as a black line. The masses of the protein-detergent-lipid complex (PDL), the detergent-lipid micelle (DL) and the protein (P) are indicated in red, blue and green, respectively.

**Figure EV3.**
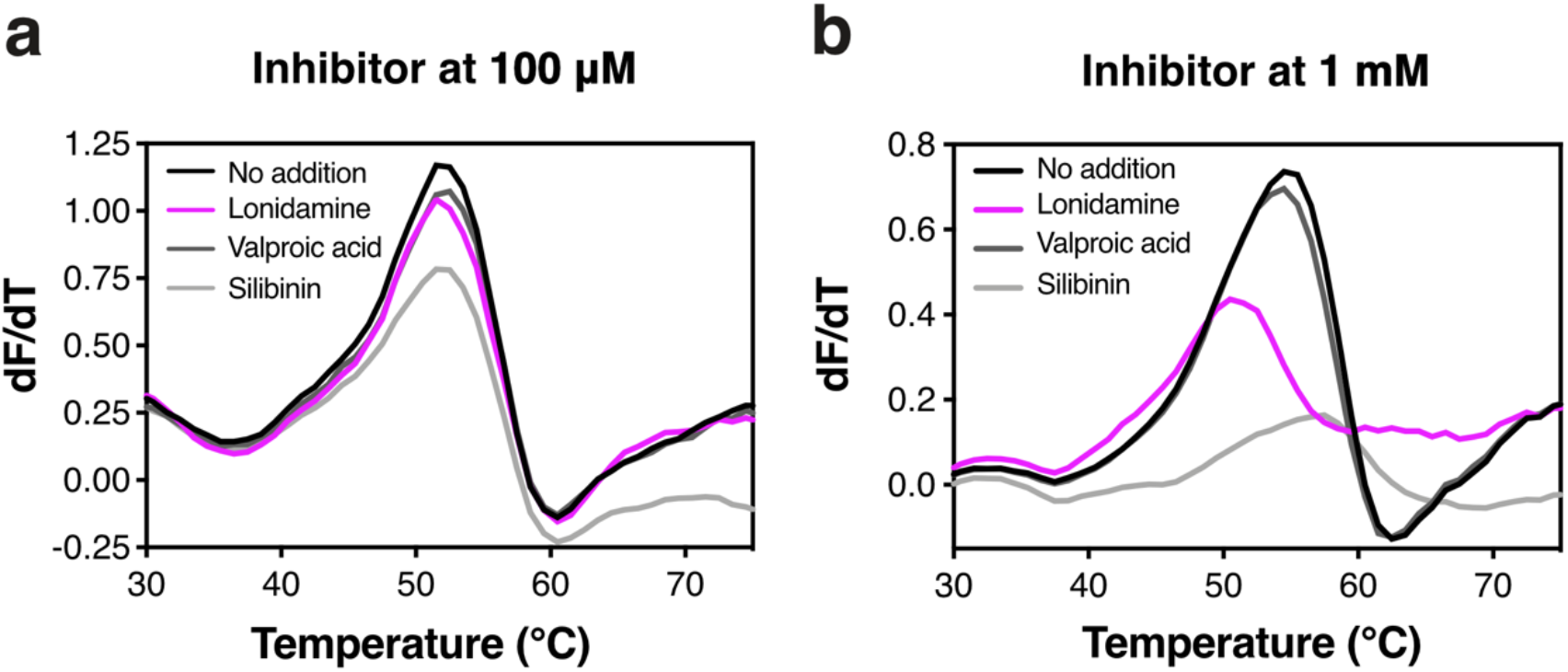
Thermostability analysis of MPC in the presence of lonidamine, silibinin and valproic acid. (**a**) First derivatives of protein unfolding curves obtained for MPC1L/MPC2 by the CPM thermostability shift assay at 100 μM, lonidamine, valproic acid or silibinin. (**b**) First derivatives of protein unfolding curves obtained for MPC1L/MPC2 by the CPM thermostability shift assay at 1 mM lonidamine, valproic acid or silibinin. Data information: The results are representative of two biological repeats, each performed in duplicate.

**Figure EV4.**
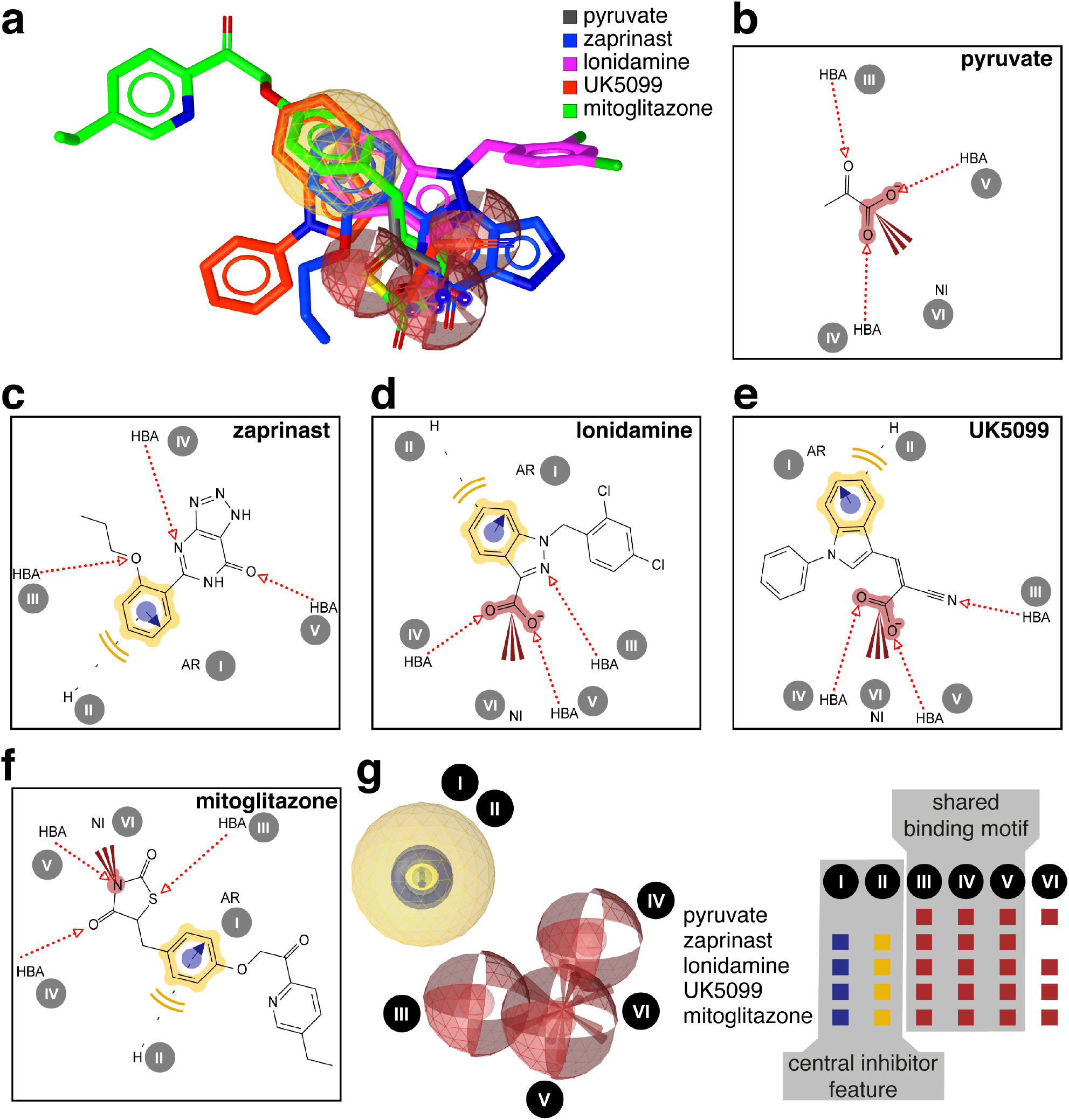
Structurally diverse MPC inhibitors have common macromolecular interactions. (**a**) Shared-feature pharmacophore model based on four prototypic MPC inhibitors and the substrate pyruvate. (**b**) Individual 2D pharmacophore representation of pyruvate. (**c**) Individual 2D pharmacophore representation zaprinast, (**d**) Individual 2D pharmacophore representation lonidamine. (**e**) Individual 2D pharmacophore representation UK5099. (**f**) Individual 2D pharmacophore representation mitoglitazone. (**g**) 3D representation of the pharmacophore with the identified features and their presence in the various prototypic MPC inhibitors. (**a-g**) Red spheres indicate hydrogen bond acceptor features, yellow spheres hydrophobic ring features, red stars indicate negative ionizable features, and blue tori indicate aromatic ring features. Pharmacophore features are abbreviated as follows: HBA, hydrogen bond acceptor; NI, negative-ionizable; H, hydrophobic ring; AR, aromatic ring. All features are identified by roman numbers.

**Figure EV5.**
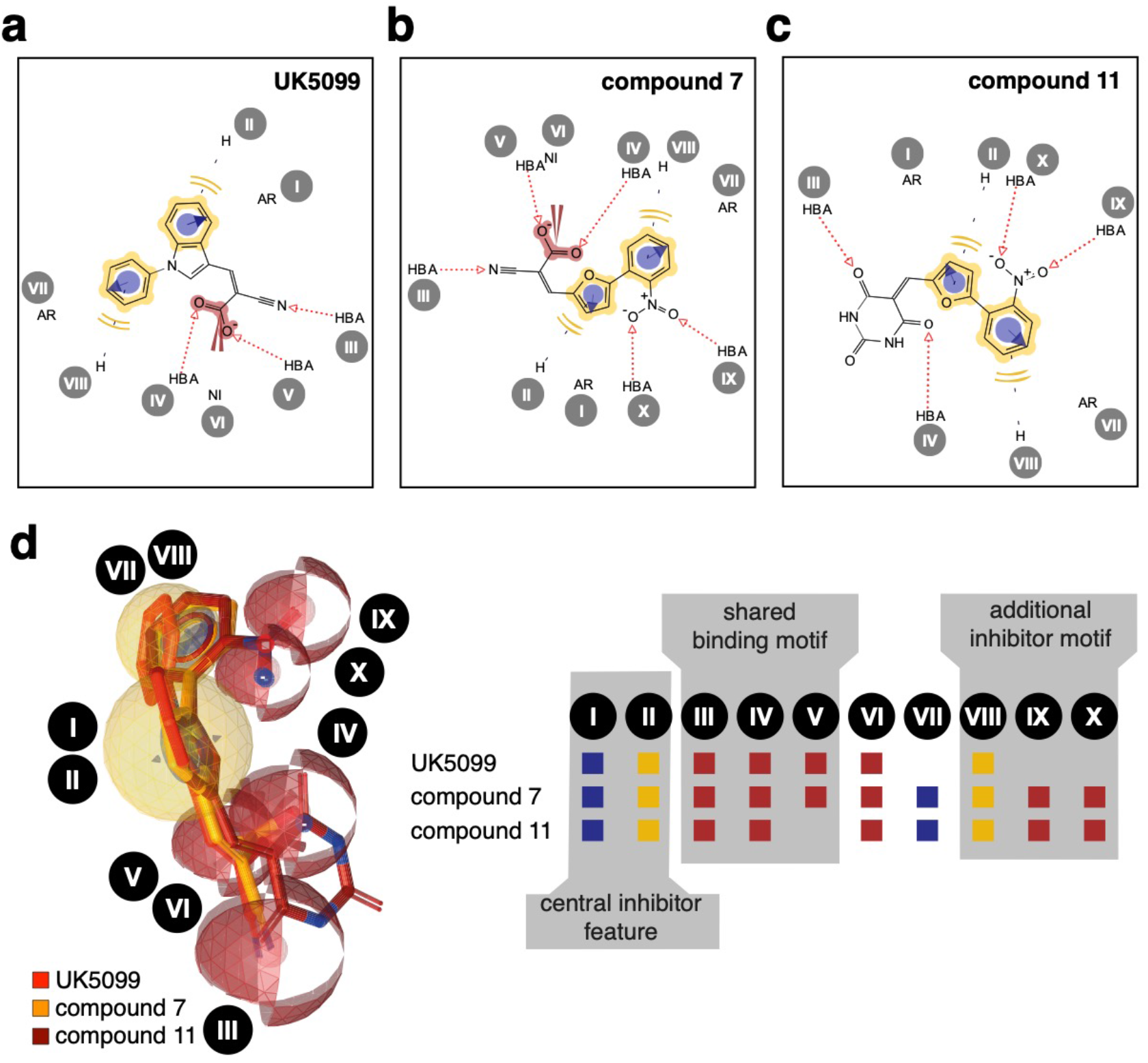
Pharmacophore matching of UK5099, compound 7 and compound 11. (**a**) 2D representation of pharmacophore properties for UK5099. (**b**) 2D representation of pharmacophore properties for compound 7. (**c**) 2D representation of pharmacophore properties for compound 11. (**d**) Shared-feature pharmacophore model based on UK5099, compound 7 and compound 11. Pharmacophore features are abbreviated as follows: HBA, hydrogen bond acceptor; NI, negative-ionizable; H, hydrophobic ring; AR, aromatic ring. All features are identified by roman numbers. Red spheres indicate hydrogen acceptor features, yellow spheres hydrophobic ring features, red stars indicate negative ionizable features, and bleu tori indicate aromatic ring features.

**Figure EV6.**
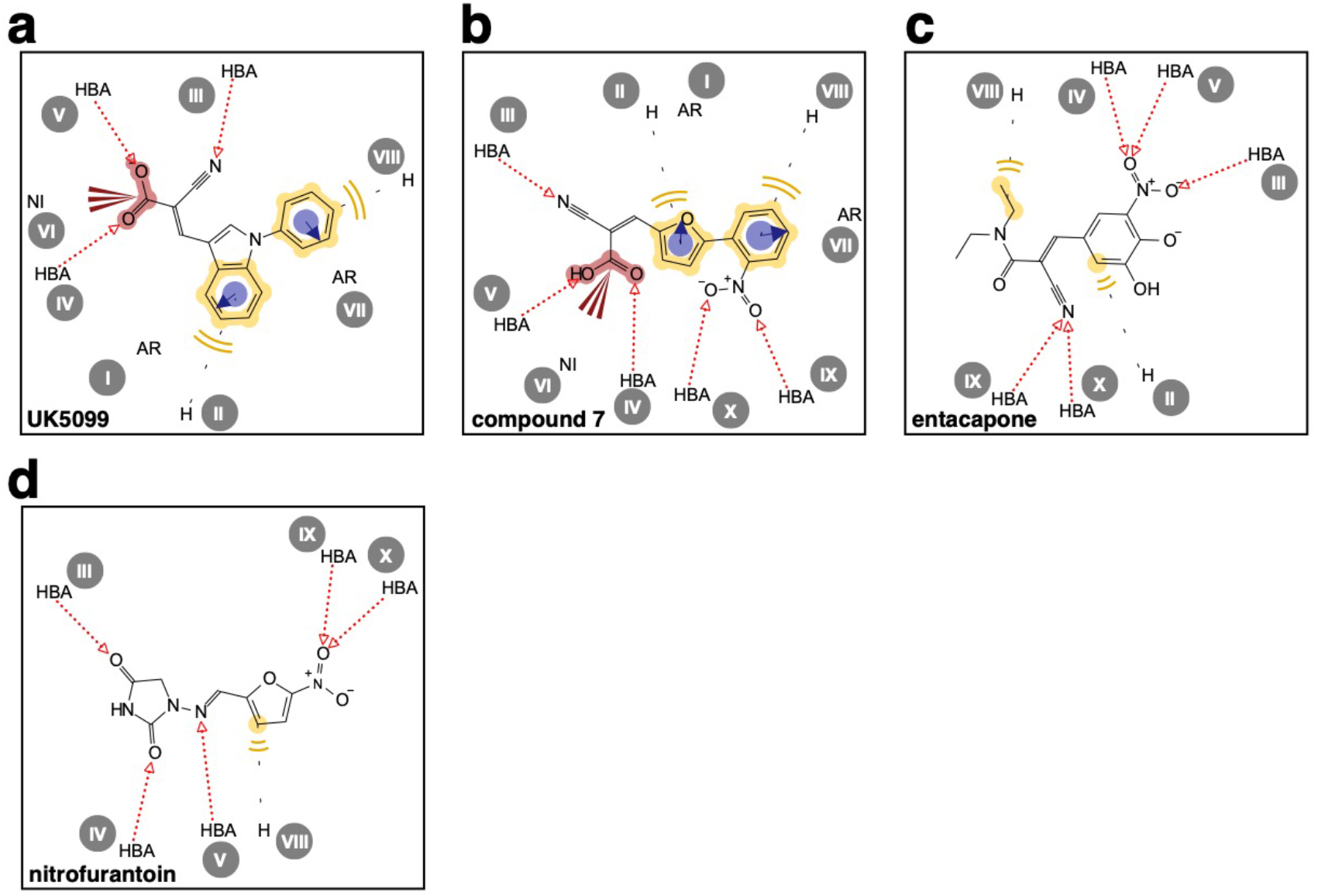
2D representation of pharmacophore properties for UK5099, compound 7, nitrofurantoin and entacapone. (**a**) 2D representation of the pharmacophore properties for UK5099. (**b**) 2D representation of the pharmacophore properties for compound 7. (**c**) 2D representation of the pharmacophore properties for entacapone. (**d**) 2D representation of the pharmacophore properties for nitrofurantoin. Pharmacophore features are abbreviated as follows: HBA, hydrogen bond acceptor; NI, negative-ionizable; H, hydrophobic ring; AR, aromatic ring.

**Table EV1.** Tm values obtained for the human MPC1L/MPC2 complex, via CPM and nano-DSF thermostability shift assays, in the presence or absence of inhibitors. Data represent the mean ± s.d. from 3 (CPM) or 2 (nano-DSF) biological repeats, each performed in duplicates or triplicates. ND stands for “not defined”.

| Compound | CPM | ΔT<sub>m</sub> | Nano-DSF | ΔT<sub>m</sub> |
| --- | --- | --- | --- | --- |
| No addition | 51.3 ± 0.2 | Not applicable | 51.9 ± 0.1 | Not applicable |
| UK5099 | 62.5 ± 0.9 | 11.2 ± 1.1 | 61.1 ± 0.6 | 9.2 ± 0.7 |
| CHC | 59.6 ± 0.4 | 8.3 ± 0.6 | 58.9 ± 0.4 | 7.0 ± 0.5 |
| Zaprinast | 58.8 ± 0.4 | 7.5 ± 0.6 | 58.2 ± 0.3 | 6.3 ± 0.4 |
| Lonidamine | 51.1 ± 0.3 | -0.2 ± 0.5 | 52.5 ± 0.2 | 0.6 ± 0.3 |
| Rosiglitazone | 56.6 ± 0.3 | 5.3 ± 0.5 | *ND* | *ND* |
| Pioglitazone | 56.4 ± 0.7 | 5.1 ± 0.9 | 53.0 ± 0.2 | 1.1 ± 0.3 |
| Mitoglitazone | 56.7 ± 0.6 | 5.4 ± 0.8 | 53.8 ± 0.4 | 1.9 ± 0.5 |
| Silibinin | 51.4 ± 0.4 | 0.1 ± 0.6 | 52.1 ± 0.2 | 0.2 ± 0.3 |
| Valproic acid | 51.3 ± 0.3 | 0.0 ± 0.5 | 51.8 ± 0.2 | -0.1 ± 0.3 |
| Entacapone | *ND* | *ND* | 58.2 ± 1.0 | 6.3 ± 1.1 |
| Nitrofurantoin | *ND* | *ND* | 54.7 ± 0.2 | 2.8 ± 0.3 |
| Compound 1 | 60.4 ± 1.0 | 9.1 ± 1.2 | *ND* | *ND* |
| Compound 2 | 63.4 ± 0.9 | 12.1 ± 1.1 | *ND* | *ND* |
| Compound 3 | 59.7 ± 1.1 | 8.4 ± 1.3 | *ND* | *ND* |
| Compound 4 | 60.4 ± 1.3 | 9.1 ± 1.5 | *ND* | *ND* |
| Compound 5 | 60.3 ± 1.4 | 9.0 ± 1.6 | *ND* | *ND* |
| Compound 6 | 59.8 ± 1.4 | 8.5 ± 1.6 | *ND* | *ND* |
| Compound 7 | 67.8 ± 1.2 | 16.5 ± 1.4 | *ND* | *ND* |
| Compound 8 | 62.3 ± 1.7 | 11.0 ± 1.9 | *ND* | *ND* |
| Compound 9 | 63.0 ± 1.8 | 11.7 ± 2.0 | *ND* | *ND* |
| Compound 10 | 51.2 ± 0.9 | -0.1 ± 1.1 | *ND* | *ND* |
| Compound 11 | 57.7 ± 2.1 | 6.4 ± 2.3 | *ND* | *ND* |
| Compound 12 | 60.4 ± 0.7 | 9.1 ± 0.9 | *ND* | *ND* |
| Compound 13 | 58.7 ± 1.7 | 7.4 ± 1.9 | *ND* | *ND* |

**Table EV2.** Results of compound preparation for pharmacophore modelling, including energy minimization using the MMFF94 protocol and generation of molecular conformations using the iCon protocol.

| Compound | No of iterations<br>energy<br>minimalization | Initial energy<br>(kcal/mol) | Minimal<br>energy<br>(kcal/mol) | No of<br>conformations |
| --- | --- | --- | --- | --- |
| Pyruvate | 116 | 85.77 | 80.66 | 1 |
| UK5099 | 226 | 192.19 | 56.70 | 14 |
| Zaprinast | 515 | 57.73 | -22.04 | 20 |
| Lonidamine | 431 | 263.29 | 128.00 | 11 |
| Mitoglitazone | 576 | 149.26 | 93.44 | 105 |
| Entacapone | 462 | 105.99 | 34.33 | 44 |
| Nitrofurantoin | 202 | 273.70 | 55.86 | 4 |
| Compound 1 | 266 | 42.36 | 10.84 | 25 |
| Compound 2 | 634 | 101.30 | 3.51 | 20 |
| Compound 7 | 532 | 132.35 | 14.78 | 16 |
| Compound 11 | 270 | 21.42 | -97.69 | 10 |

## Acknowledgments

We thank Professor Jean-Claude Martinou for kindly providing the SHY15 *mpc* deletion strain. We also thank Dr. Chris Johnson for access to the NanoTemper Prometheus NT.48 at the MRC Laboratory of Molecular Biology, Dr. Shane Palmer for the large-scale fermentation at the Mitochondrial Biology Unit and Dr. Caterina Garone for advice on small molecule drugs associated with mitochondrial toxicity. This work was funded by the Medical Research Council UK (Grant MC_UU_00015/1). T.J.J.S. was supported by a Long-Term EMBO Fellowship of the European Molecular Biology Organization ALTF 268-2016.

## Contributions

Conception and design of the research: S.T., T.J.J.S., and E.R.S.K., Molecular Biology: S.T., D.T.D.J., M.S.K., Biochemical and Biophysical data acquisition: S.T., V.M., C.T., Pharmacophore modelling: T.J.J.S., Mass Spectrometry: S.D., and I.M.F., Data analysis: S.T. and T.J.J.S., Data interpretation: S.T., T.J.J.S., I.M.F, and E.R.S.K., Writing of the paper: S.T., T.J.J.S. and E.R.S.K. All authors read and revised the paper.

## REFERENCES

Bader DA, Hartig SM, Putluri V, Foley C, Hamilton MP, Smith EA, Saha PK, Panigrahi A, Walker C, Zong L, Martini-Stoica H, Chen R, Rajapakshe K, Coarfa C, Sreekumar A, Mitsiades N, Bankson JA, Ittmann MM, O’Malley BW, Putluri N et al. (2019) Mitochondrial pyruvate import is a metabolic vulnerability in androgen receptor-driven prostate cancer. Nat Metab 1: 70–85

Benavides J, Martin A, Ugarte M, Valdivieso F (1982) Inhibition by valproic acid of pyruvate uptake by brain mitochondria. Biochem Pharmacol 31: 1633–1636

Bender T, Pena G, Martinou JC (2015) Regulation of mitochondrial pyruvate uptake by alternative pyruvate carrier complexes. EMBO J 34: 911–924

Bricker DK, Taylor EB, Schell JC, Orsak T, Boutron A, Chen YC, Cox JE, Cardon CM, Van Vranken JG, Dephoure N, Redin C, Boudina S, Gygi SP, Brivet M, Thummel CS, Rutter J (2012) A mitochondrial pyruvate carrier required for pyruvate uptake in yeast, Drosophila, and humans. Science 337: 96–100

Buchan DW, Minneci F, Nugent TC, Bryson K, Jones DT (2013) Scalable web services for the PSIPRED Protein Analysis Workbench. Nucleic Acids Res 41: W349–357

Carbonera D, Angrilli A, Azzone GF (1988) Mechanism of nitrofurantoin toxicity and oxidative stress in mitochondria. Biochim Biophys Acta 936: 139–147

Colca JR, McDonald WG, Adams WJ (2018) MSDC-0602K, a metabolic modulator directed at the core pathology of non-alcoholic steatohepatitis. Expert Opin Investig Drugs 27: 631–636

Colca JR, McDonald WG, Cavey GS, Cole SL, Holewa DD, Brightwell-Conrad AS, Wolfe CL, Wheeler JS, Coulter KR, Kilkuskie PM, Gracheva E, Korshunova Y, Trusgnich M, Karr R, Wiley SE, Divakaruni AS, Murphy AN, Vigueira PA, Finck BN, Kletzien RF (2013) Identification of a mitochondrial target of thiazolidinedione insulin sensitizers (mTOT)--relationship to newly identified mitochondrial pyruvate carrier proteins. PLoS One 8: e61551

Colca JR, Tanis SP, McDonald WG, Kletzien RF (2014) Insulin sensitizers in 2013: new insights for the development of novel therapeutic agents to treat metabolic diseases. Expert Opin Investig Drugs 23: 1–7

Colturato CP, Constantin RP, Maeda AS, Jr., Constantin RP, Yamamoto NS, Bracht A, Ishii-Iwamoto EL, Constantin J (2012) Metabolic effects of silibinin in the rat liver. Chem Biol Interact 195: 119–132

Compan V, Pierredon S, Vanderperre B, Krznar P, Marchiq I, Zamboni N, Pouyssegur J, Martinou JC (2015) Monitoring Mitochondrial Pyruvate Carrier Activity in Real Time Using a BRET-Based Biosensor: Investigation of the Warburg Effect. Mol Cell 59: 491–501

Corbet C, Bastien E, Draoui N, Doix B, Mignion L, Jordan BF, Marchand A, Vanherck JC, Chaltin P, Schakman O, Becker HM, Riant O, Feron O (2018) Interruption of lactate uptake by inhibiting mitochondrial pyruvate transport unravels direct antitumor and radiosensitizing effects. Nat Commun 9: 1208

Crichton PG, Lee Y, Ruprecht JJ, Cerson E, Thangaratnarajah C, King MS, Kunji ER (2015) Trends in thermostability provide information on the nature of substrate, inhibitor, and lipid interactions with mitochondrial carriers. J Biol Chem 290: 8206–8217

Divakaruni AS, Wallace M, Buren C, Martyniuk K, Andreyev AY, Li E, Fields JA, Cordes T, Reynolds IJ, Bloodgood BL, Raymond LA, Metallo CM, Murphy AN (2017) Inhibition of the mitochondrial pyruvate carrier protects from excitotoxic neuronal death. J Cell Biol 216: 1091–1105

Divakaruni AS, Wiley SE, Rogers GW, Andreyev AY, Petrosyan S, Loviscach M, Wall EA, Yadava N, Heuck AP, Ferrick DA, Henry RR, McDonald WG, Colca JR, Simon MI, Ciaraldi TP, Murphy AN (2013) Thiazolidinediones are acute, specific inhibitors of the mitochondrial pyruvate carrier. Proc Natl Acad Sci U S A 110: 5422–5427

Du J, Cleghorn WM, Contreras L, Lindsay K, Rountree AM, Chertov AO, Turner SJ, Sahaboglu A, Linton J, Sadilek M, Satrustegui J, Sweet IR, Paquet-Durand F, Hurley JB (2013) Inhibition of mitochondrial pyruvate transport by zaprinast causes massive accumulation of aspartate at the expense of glutamate in the retina. J Biol Chem 288: 36129–36140

Elia I, Rossi M, Stegen S, Broekaert D, Doglioni G, van Gorsel M, Boon R, Escalona-Noguero C, Torrekens S, Verfaillie C, Verbeken E, Carmeliet G, Fendt SM (2019) Breast cancer cells rely on environmental pyruvate to shape the metastatic niche. Nature 568: 117–121

Gasteiger E, Hoogland C, Gattiker A, Duvaud Se, Wilkins MR, Appel RD, Bairoch A (2005) Protein Identification and Analysis Tools on the ExPASy Server. In The Proteomics Protocols Handbook, Walker JM (ed) pp 571–607. Humana Press

Ghosh A, Tyson T, George S, Hildebrandt EN, Steiner JA, Madaj Z, Schulz E, Machiela E, McDonald WG, Escobar Galvis ML, Kordower JH, Van Raamsdonk JM, Colca JR, Brundin P (2016) Mitochondrial pyruvate carrier regulates autophagy, inflammation, and neurodegeneration in experimental models of Parkinson’s disease. Sci Transl Med 8: 368ra174

Gietz RD, Schiestl RH (2007) High-efficiency yeast transformation using the LiAc/SS carrier DNA/PEG method. Nat Protoc 2: 31–34

Gray LR, Sultana MR, Rauckhorst AJ, Oonthonpan L, Tompkins SC, Sharma A, Fu X, Miao R, Pewa AD, Brown KS, Lane EE, Dohlman A, Zepeda-Orozco D, Xie J, Rutter J, Norris AW, Cox JE, Burgess SC, Potthoff MJ, Taylor EB (2015) Hepatic Mitochondrial Pyruvate Carrier 1 Is Required for Efficient Regulation of Gluconeogenesis and Whole-Body Glucose Homeostasis. Cell Metab 22: 669–681

Grunig D, Felser A, Bouitbir J, Krahenbuhl S (2017) The catechol-O-methyltransferase inhibitors tolcapone and entacapone uncouple and inhibit the mitochondrial respiratory chain in HepaRG cells. Toxicol In Vitro 42: 337–347

Grunig D, Felser A, Duthaler U, Bouitbir J, Krahenbuhl S (2018) Effect of the Catechol-O-Methyltransferase Inhibitors Tolcapone and Entacapone on Fatty Acid Metabolism in HepaRG Cells. Toxicol Sci 164: 477–488

Halestrap AP (1975) The mitochondrial pyruvate carrier. Kinetics and specificity for substrates and inhibitors. Biochem J 148: 85–96

Halestrap AP (1976) The mechanism of the inhibition of the mitochondrial pyruvate transportater by alpha-cyanocinnamate derivatives. Biochem J 156: 181–183

Halestrap AP, Denton RM (1974) Specific inhibition of pyruvate transport in rat liver mitochondria and human erythrocytes by alpha-cyano-4-hydroxycinnamate. Biochem J 138: 313–316

Halestrap AP, Denton RM (1975) The specificity and metabolic implications of the inhibition of pyruvate transport in isolated mitochondria and intact tissue preparations by alpha-Cyano-4-hydroxycinnamate and related compounds. Biochem J 148: 97–106

Harrison SA, Alkhouri N, Davison BA, Sanyal A, Edwards C, Colca JR, Lee BH, Loomba R, Cusi K, Kolterman O, Cotter G, Dittrich HC (2020) Insulin sensitizer MSDC-0602K in non-alcoholic steatohepatitis: A randomized, double-blind, placebo-controlled phase IIb study. J Hepatol 72: 613–626

Herzig S, Raemy E, Montessuit S, Veuthey JL, Zamboni N, Westermann B, Kunji ER, Martinou JC (2012) Identification and functional expression of the mitochondrial pyruvate carrier. Science 337: 93–96

Jones DT, Taylor WR, Thornton JM (1994) A model recognition approach to the prediction of all-helical membrane protein structure and topology. Biochemistry 33: 3038–3049

Lee J, Jin Z, Lee D, Yun JH, Lee W (2020) Characteristic Analysis of Homo- and Heterodimeric Complexes of Human Mitochondrial Pyruvate Carrier Related to Metabolic Diseases. Int J Mol Sci 21

McCommis KS, Hodges WT, Brunt EM, Nalbantoglu I, McDonald WG, Holley C, Fujiwara H, Schaffer JE, Colca JR, Finck BN (2017) Targeting the mitochondrial pyruvate carrier attenuates fibrosis in a mouse model of nonalcoholic steatohepatitis. Hepatology 65: 1543–1556

Miller CA, 3rd, Martinat MA, Hyman LE (1998) Assessment of aryl hydrocarbon receptor complex interactions using pBEVY plasmids: expressionvectors with bi-directional promoters for use in Saccharomyces cerevisiae. Nucleic Acids Res 26: 3577–3583

Nagampalli RSK, Quesnay JEN, Adamoski D, Islam Z, Birch J, Sebinelli HG, Girard R, Ascencao CFR, Fala AM, Pauletti BA, Consonni SR, de Oliveira JF, Silva ACT, Franchini KG, Leme AFP, Silber AM, Ciancaglini P, Moraes I, Dias SMG, Ambrosio ALB (2018) Human mitochondrial pyruvate carrier 2 as an autonomous membrane transporter. Sci Rep 8: 3510

Nancolas B, Guo L, Zhou R, Nath K, Nelson DS, Leeper DB, Blair IA, Glickson JD, Halestrap AP (2016) The anti-tumour agent lonidamine is a potent inhibitor of the mitochondrial pyruvate carrier and plasma membrane monocarboxylate transporters. Biochem J 473: 929–936

Nanjan MJ, Mohammed M, Prashantha Kumar BR, Chandrasekar MJN (2018) Thiazolidinediones as antidiabetic agents: A critical review. Bioorg Chem 77: 548–567

Nath K, Guo L, Nancolas B, Nelson DS, Shestov AA, Lee SC, Roman J, Zhou R, Leeper DB, Halestrap AP, Blair IA, Glickson JD (2016) Mechanism of antineoplastic activity of lonidamine. Biochim Biophys Acta 1866: 151–162

Papa S, Paradies G (1974) On the mechanism of translocation of pyruvate and other monocarboxylic acids in rat-liver mitochondria. Eur J Biochem 49: 265–274

Paradies G, Papa S (1975) The transport of monocarboxylic oxoacids in rat liver mitochondria. FEBS Lett 52: 149–152

Quansah E, Peelaerts W, Langston JW, Simon DK, Colca J, Brundin P (2018) Targeting energy metabolism via the mitochondrial pyruvate carrier as a novel approach to attenuate neurodegeneration. Mol Neurodegener 13: 28

Shah RC, Matthews DC, Andrews RD, Capuano AW, Fleischman DA, VanderLugt JT, Colca JR (2014) An evaluation of MSDC-0160, a prototype mTOT modulating insulin sensitizer, in patients with mild Alzheimer’s disease. Curr Alzheimer Res 11: 564–573

Sievers F, Wilm A, Dineen D, Gibson TJ, Karplus K, Li W, Lopez R, McWilliam H, Remmert M, Soding J, Thompson JD, Higgins DG (2011) Fast, scalable generation of high-quality protein multiple sequence alignments using Clustal Omega. Mol Syst Biol 7: 539

Slotboom DJ, Duurkens RH, Olieman K, Erkens GB (2008) Static light scattering to characterize membrane proteins in detergent solution. Methods 46: 73–82

Soccio RE, Chen ER, Lazar MA (2014) Thiazolidinediones and the promise of insulin sensitization in type 2 diabetes. Cell Metab 20: 573–591

Tavoulari S, Thangaratnarajah C, Mavridou V, Harbour ME, Martinou JC, Kunji ER (2019) The yeast mitochondrial pyruvate carrier is a hetero-dimer in its functional state. EMBO J 38

Thangaratnarajah C, Ruprecht JJ, Kunji ERS (2014) Calcium-induced conformational changes of the regulatory domain of human mitochondrial aspartate/glutamate carriers. Nat Commun 5: 5491

Tompkins SC, Sheldon RD, Rauckhorst AJ, Noterman MF, Solst SR, Buchanan JL, Mapuskar KA, Pewa AD, Gray LR, Oonthonpan L, Sharma A, Scerbo DA, Dupuy AJ, Spitz DR, Taylor EB (2019) Disrupting Mitochondrial Pyruvate Uptake Directs Glutamine into the TCA Cycle away from Glutathione Synthesis and Impairs Hepatocellular Tumorigenesis. Cell Rep 28: 2608–2619 e2606

Vanderperre B, Cermakova K, Escoffier J, Kaba M, Bender T, Nef S, Martinou JC (2016) MPC1-like Is a Placental Mammal-specific Mitochondrial Pyruvate Carrier Subunit Expressed in Postmeiotic Male Germ Cells. J Biol Chem 291: 16448–16461

Wilcken R, Zimmermann MO, Lange A, Joerger AC, Boeckler FM (2013) Principles and applications of halogen bonding in medicinal chemistry and chemical biology. J Med Chem 56: 1363–1388

Yamashita Y, Vinogradova EV, Zhang X, Suciu RM, Cravatt BF (2020) A Chemical Proteomic Probe for the Mitochondrial Pyruvate Carrier Complex. Angew Chem Int Ed Engl 59: 3896–3899

